# The neuroreceptors and transporters underlying spontaneous brain activity

**DOI:** 10.1101/2024.12.06.627265

**Authors:** Johan Nakuci, Kanika Bansal

**Affiliations:** U.S. ARMY DEVCOM Army Research Laboratory, Aberdeen Proving Ground, Aberdeen, MD, 21005, USA; Computer Science and Electrical Engineering, University of Maryland Baltimore County, Baltimore, MD, 21250, USA

**Keywords:** molecular-fMRI, resting-state fMRl, neuroreceptors, neuropsychiatric illness, LSD, Modafinil, structural and cortical geometric modes

## Abstract

Determining the neuromodulators driving brain activity is critical for understanding cognition and neuropathology. Neuromodulators act through neuroreceptors in a coordinated manner, yet their interconnected dynamics are often overlooked. We show that a neuroreceptor-based modeling framework using cortical density maps of 19 neuroreceptors and transporters from Positron Emission Tomography (PET) can model BOLD-derived brain activity. This framework reconstructs activity across four datasets (N = 314) identifying two neuroreceptor modules linked to higher-order associative or somatomotor and visual networks. Applying the framework to independent datasets, we recover the binding profiles of LSD and Modafinil, demonstrating consistency with known pharmacological and neurobiological associations. Additionally, it uncovered associations between neuroreceptors, transporters, and altered brain activity in neuropsychiatric disorders. These findings demonstrate the framework’s potential to elucidate neuromodulatory mechanisms and advance our understanding of brain function across diverse states and conditions.

## Introduction

At any given moment, across the brain, a diverse array of neuromodulators binds to and unbinds from neurons driving activity and shaping the large-scale brain dynamics. The complexity of this process becomes evident when considering that the human brain utilizes over 100 neuromodulators across approximately 10^11^ neurons, each forming an average of 1,000 unique connections^1^. This intricate interplay presents an opportunity, albeit with a significant challenge, to understand how the interactions of specific neuromodulators drive the large-scale brain activity in healthy brains and how these interactions change under specific pathological conditions.

Neuromodulators operate through diverse neuroreceptor subtypes with varying signaling cascades^2^, spatial distributions^3^, and temporal dynamics^4^ to maintain the excitation/inhibition balance enabling the brain’s vast computational capabilities^5^. These dynamic interactions underly how molecular-level receptor activity translates to systems-level effects. For instance, tonic and bursting activity of the locus coeruleus which releases norepinephrine can shift network activity across the cortical hierarchy modulating multiple brain states^4^. Moreover, the balance between excitatory and inhibitory inputs dynamically changes over time, reflecting different brain states^6^. These fluctuations have been linked to aging^7^, psychosis^8^ and can impact cognitive processes and behaviors across diverse domains, including social interactions^9^, economic decisions^10^, and even perception^11^.

However, most studies investigate the relationship between one or two neuromodulators and brain activity at a time. Partly this is due to limitations of current methods as they typically measure only a few neuromodulators at a time or are restricted to a subset of brain regions^12,13^. Crucially, an understanding of the dynamic relationship among multiple neuromodulators that drive large-scale brain activity is lacking.

Building upon the burgeoning field of molecular-fMRI^14^ we present a neuroreceptor-based framework to unravel the dynamic relationship among multiple neuroreceptors and transporters underlying spontaneous brain activity. Previous work has investigated these questions utilizing spatial correlation through neuroreceptor informed network control^15,16^ or whole brain modelling^17^. Collectively, these approaches have the potential to unravel the link between molecular-level neuroreceptor activity and systems-level effects^18,19^.

The concentration of neuroreceptors is an important factor in mediating the effects of neuromodulators on brain activity, as these serve as the primary sites for excitatory synaptic inputs^20^, dynamically changing in response to neural activity, stimuli and conditions^21^, and can shape long-term memory^22^. The concentration of neuroreceptors across the brain can be non-invasively estimated using PET tracer imaging^23,24^. Moreover, recent work has shown that neuroreceptors shape brain-wide communication estimated from the BOLD signal^15,25,26^. This association suggests that combining neuroreceptor density maps estimated from PET imaging with the BOLD signal could be a non-invasive method to uncover the neuromodulators that drive brain activity across the human cortex and unravel the complex mechanisms that underlie cognition, behavior and neuropathology.

Here we show that a neuroreceptor-based framework can reconstruct the BOLD signal through a weighted summation of PET-derived neuroreceptor and transporter density maps. Leveraging this framework, we reveal that two distinct modules of neuroreceptors and transporters linked to higher-order associative or visual and somatomotor networks underly the BOLD signal. Moreover, the framework can recover neuroreceptors and transporters — 5-HT1a, 5-HT1b, 5-HT2a, and D2 — known to bind lysergic acid diethylamide (LSD), and D2, 5-HT1a, 5-HT2a, and NET mediating the effects of Modafinil demonstrating consistency with known pharmacological and neurobiological associations. Further, we demonstrate the framework’s utility by identifying associations linking neuroreceptors, transporters, and altered brain activity in schizophrenia, bipolar disorder, and ADHD.

## Results

### Reconstructing the BOLD signal and functional connectivity from neuroreceptor and transporter maps

To determine if neuroreceptor and transporter density maps could non-invasively reveal the neuromodulators that drive spontaneous brain activity, we assessed whether a linear modeling framework could reconstruct the temporal dynamics of the BOLD signal. We utilize the most comprehensive neuroreceptors and transporters cortical density maps to date as detailed in Hansen et. al 2022^27^. In brief, PET tracer imaging from a combined 1,238 healthy individuals was used to map the densities of 19 neuroreceptors and transporters from 9 neuromodulator systems: dopamine (D1, D2, DAT), norepinephrine (NET), serotonin (5-HT1A, 5-HT1B, 5-HT2A, 5-HT4, 5-HT6, 5-HTT), acetylcholine (α4β2, M1, VAChT), glutamate (mGluR5, NMDA), γ-aminobutyric acid (GABAa), histamine (H3), cannabinoid (CB1), and opioid neuroreceptor (MOR; **Fig. S1**). For each neuroreceptor and transporter, we estimated the average density within 200 brain regions based on the local-global parcellation of the human cerebral cortex estimated from resting-state fMRI by Schaefer et. al., 2018 because voxel-wise estimates can be unstable and noisy^28^.

Spontaneous brain activity was estimated from 15 minutes of resting-state fMRI (N = 48) part of the Human Connectome Project (HCP)^29^. The extensive scanning duration of the HCP dataset allowed for an estimation of the relationship between neuroreceptors, transporters, and brain activity across a broader range of neural states. The voxelwise BOLD signal from each participant was averaged across the same brain regions defined by the Schaefer atlas, resulting in 200 x 1,200 data points per participant and 200 x 57,600 data points across all participants. Independently, each of the 1,200 time points of the BOLD signal were reconstructed from a weighted linear summation of the 19 neuroreceptor and transporter density maps (**Fig. 1A**). To quantify effectiveness of the neuroreceptor-based framework, Pearson’s correlation was used to estimate the accuracy between the reconstructed and the empirical brain maps for each of the 57,600 time points.

**Figure 1.**
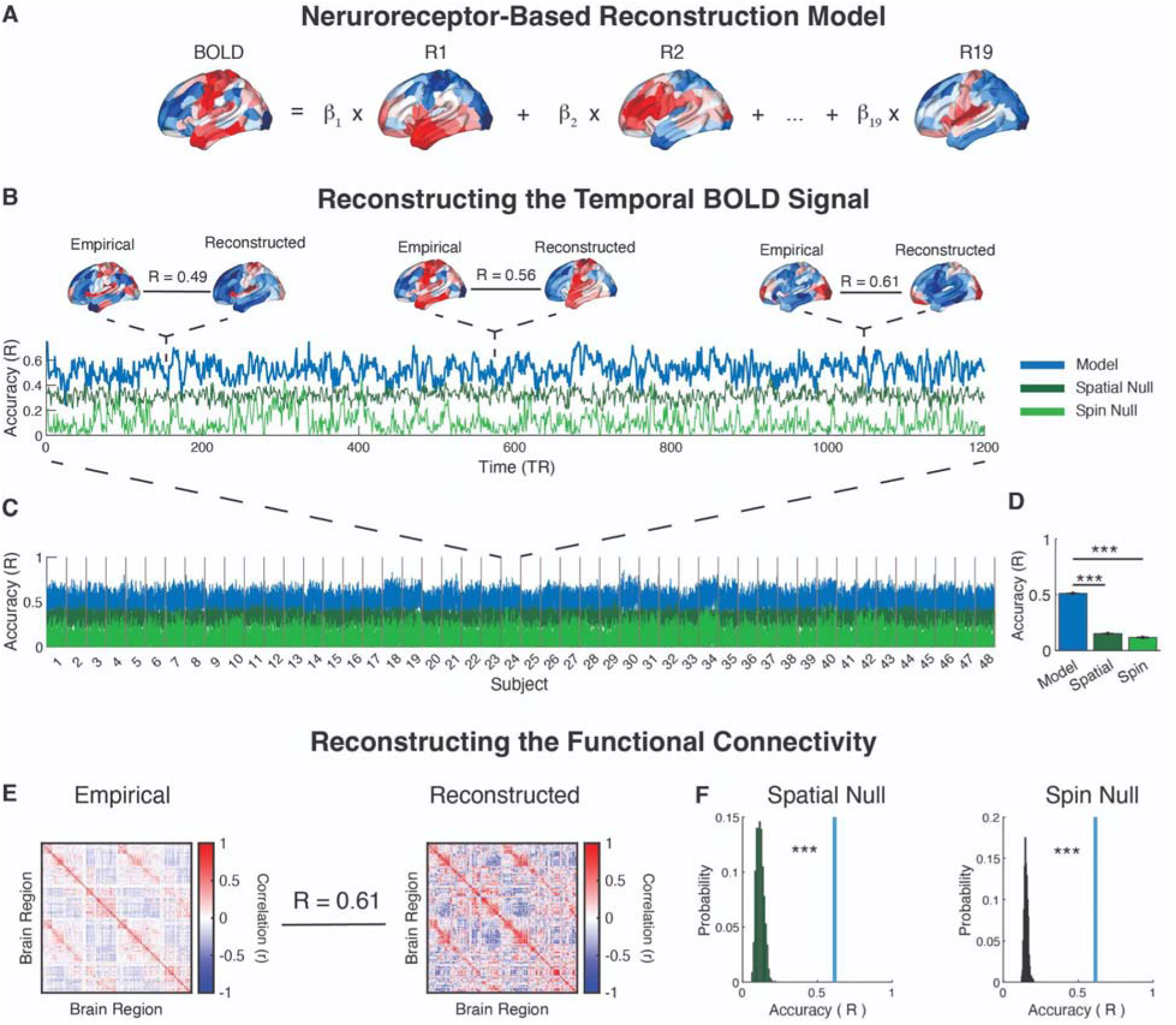
Neuroreceptor-based framework can reconstruct the BOLD signal and functional connectivity. (A) Neuroreceptor-based framework schematic. The BOLD signal was reconstructed from a linear summation of 19 neuroreceptors and transporters (R_1_ to R_19_) each weighted by corresponding coefficients (β_1_ to β_19_). For visualization purposes we have omitted the constant term in the model. (B) Reconstruction accuracy of the BOLD signal. Accuracy of neuroreceptor-based framework in reconstructing the BOLD signal over time in a representative participant. Pearson’s correlation was used to estimate the accuracy between the reconstructed and the empirical brain maps. Brain plots depict the empirical, reconstructed brain and the accuracy between brain maps for representative time points. Line plots depict accuracy over time. Blue line reflects the observed accuracy of the neuroreceptor-based model. Dark and light green lines reflect the accuracy from spatially permuted (spatial) and spin-permutation (spin) null models. (C) Group level reconstruction accuracy for each of the 1,200 time points per participant. (D) Neuroreceptor-based reconstruction is significantly greater than spatial and spin null models. Bar plots show the average accuracy at the group-level and for the spatial and spin null models. Error bars show mean ± sem. (E) Reconstructing functional connectivity. Accuracy of reconstructing the group-level functional connectivity. The BOLD signal was concatenated across participants and Pearson’s correlation was used to estimate the empirical functional connectivity (*left*). The reconstructed BOLD signal was concatenated across participants and Pearson’s correlation was used to estimate the functional connectivity (*right*). (F) Comparison with spatial and spin null models. Blue line depicts the accuracy value from panel D. For both null models, the functional connectivity was estimated from the reconstructed BOLD signal by first concatenating the signal across participants followed by Pearson’s correlation. ***P < 0.001.

Focusing on a representative participant, the BOLD signal at each moment in time was reconstructed with an average accuracy of R_BOLD_subj24_ = 0.53 ± 0.08 (mean ± std; **Fig. 1B**). Comparable accuracy was observed across all participants (R_BOLD_ = 0.51 ± 0.08; **Fig. 1C**). The observed reconstruction accuracy was significantly greater than that obtained from null models based on spatially permuted neuroreceptor and transporter maps (P_spatial_ < 0.001) and from a permutation that preserved the spatial auto-correlation, spin test (P_spin_ < 0.001; **Fig. 1D**)^25^. Moreover, the accuracy in reconstruction was differentially distributed across the brain with the visual network exhibiting on average higher accuracy (**Fig. S2**).

Further, given that spontaneous brain activity exhibits coordination, or functional connectivity (FC), among brain regions, we determined whether the reconstructed BOLD signal preserved this inherent property. Moreover, reconstructing the BOLD signal from neuroreceptor and transporter density does not inherently guarantee that the reconstruction will preserve FC at a comparable level. Thus, the ability to reconstruct the inherent FC would add to the reliability of the results. Conversely, if the reconstruction were dominated by random noise, the inherent FC would not be reproduced.

To assess whether the reconstructed BOLD signal preserved inherent functional connectivity, we concatenated the empirical and reconstructed BOLD signals across participants and used Pearson’s correlation to estimate the empirical and reconstructed functional connectivity networks, respectively. The reconstructed functional connectivity demonstrated comparable accuracy to that observed in the temporal BOLD signal (R_FC_ = 0.61; **Fig. 1E**) and was significantly greater than both null models (P_spatial_ < 0.001; P_spin_ < 0.001; **Fig. 1F**). Together, these findings highlight that a neuroreceptor-based framework can reconstruct spontaneous brain activity and its inherent functional connectivity.

### Two distinct modules of neuroreceptors and transporters underlie dynamic brain activity

No neuroreceptor or transporter acts in isolation; their actions are inherently coordinated^26^. However, most studies are limited to investigating only a few neuroreceptors and transporters at a time and therefore a comprehensive view is crucially lacking. To address this lack of knowledge, we investigated if the neuroreceptor-based framework could unravel the coordinated interplay among neuroreceptors and transporters working together to drive brain activity.

As shown in Figure 2A, these drive values (i.e. β-values) exhibit marked fluctuations over time within a participant. These observations may suggest that fluctuations in brain activity reflects coordination among multiple and distinct neuroreceptors and transporters (**Fig. 2B**). Understanding how neuromodulatory systems influence the emergent patterns of brain activity and cognitive processes requires quantifying the extent of temporal coordination among neuroreceptors and transporters.

**Figure 2.**
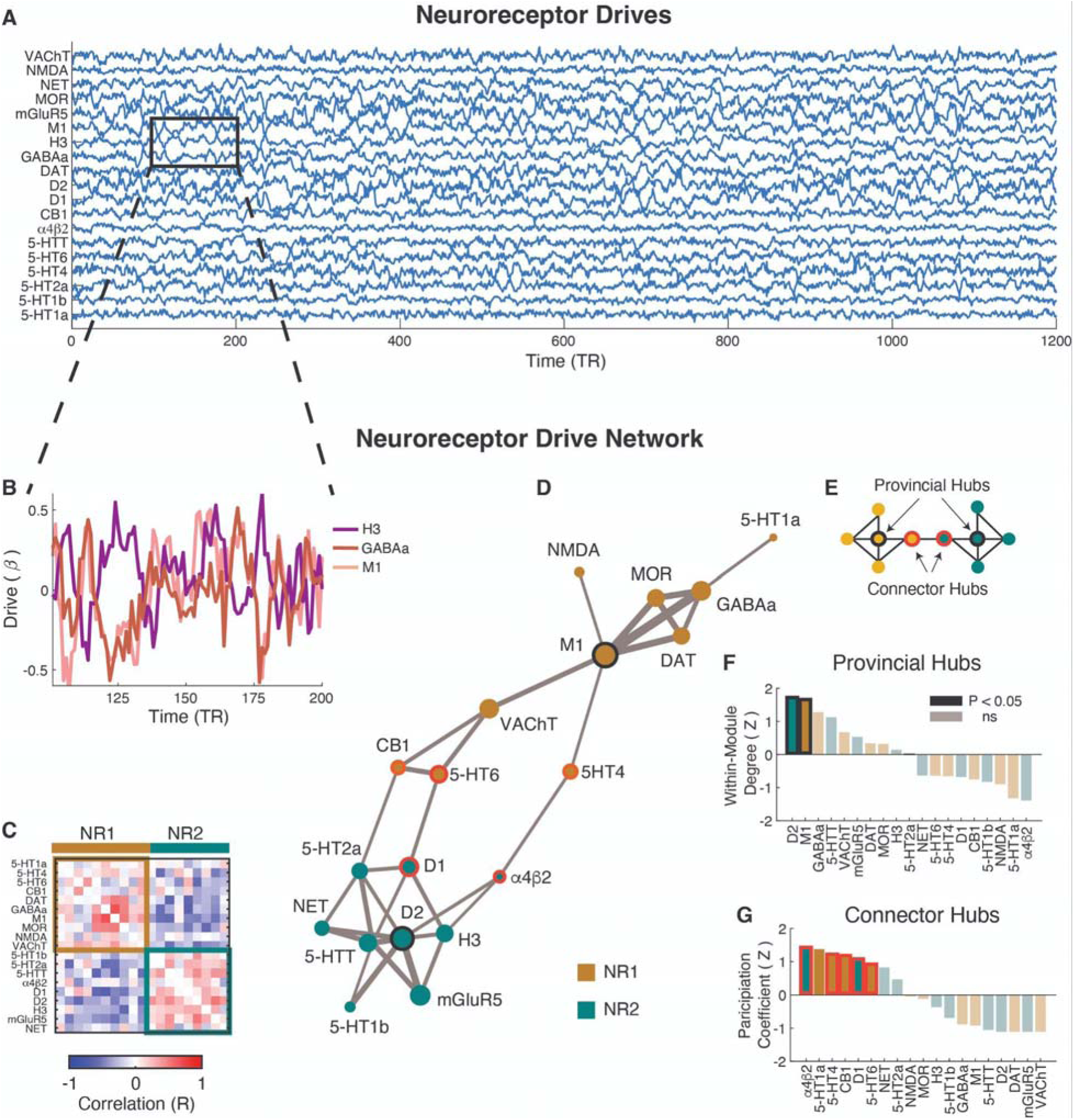
Two neuroreceptor and transporter modules underlie spontaneous brain activity. (A) Drive values (β-values) for all 19 neuroreceptors and transporters from a representative participant. (B) Drive values for GABAa, M1, and H3 neuroreceptors over a period of time from the same participant. (C) Group-level drive network depicting the strength of coordination between neuroreceptor drive values over time. Drive network was estimated using Pearson correlation to quantify the extent of pairwise coordination among neuroreceptors and transporters over time. Clustering analysis identified two distinct modules, NR1 and NR2. (D) Visualization of the drive network structure, showing only the top 20% of strongest connections. Node size indicates the weighted degree of the network, while edge thickness reflects the strength of connections (R-value) between neuroreceptors. The black outline represents provincial hubs, and the red outline represents connector hubs. (E) Schematic representation of provincial and connector hubs in a network. (F) Provincial hub and (G) connector hub scores for each neuroreceptor, ranked according to hub score. Hub scores were calculated using positive values in the neuroreceptor drive network, and statistical significance was assessed by permuting the drive network in Panel C 1,000 times. Outlined bars represent identified hubs. Note that while 5-HT1a exhibited a high connector hub score, it was not considered a hub due to weak correlation values within and between modules. ns, non-significant.

To quantify the extent of coordination among neuroreceptors and transporters, the drive values were concatenated across participants and pairwise coordination was estimated using Pearson correlation. Positive correlations indicated coordinated changes, as observed between GABAa and M1 (R_GABAa:M1_ = 0.68), while negative correlations suggested inverse relationships, such as between GABAa and H3 (R_GABAa:H3_ = -0.29). Across all neuroreceptors and transporters, the correlations form a neuroreceptor drive network representing the overall coordination among all neuroreceptors and transporters.

The neuroreceptor drive network revealed distinct patterns of positive and negative correlations among drive values, consistent with those observed between GABAa, M1, and H3 (**Fig. 2C,D**). Moreover, clustering analysis identified two modules, NR1 and NR2 of neuroreceptors and transporters. Additionally, as shown in Figure 2D, a subset of neuroreceptors within both the NR1 and NR2 modules exhibited strong coordination with other neuroreceptors in their respective modules (black outlined nodes), while another subset acted as connectors linking the NR1 and NR2 modules (red outlined nodes).

Neuroreceptors or transporters that showed strong coordination (i.e., positive correlations) within their module are referred to as provincial hubs, whereas those linking different modules are termed connector hubs (**Fig. 2E**). To quantify the role of a neuroreceptor or transporter as a provincial hub, we calculated the within-module degree, which measures the strength of coordination within the same module. The M1 neuroreceptor was identified as a candidate provincial hub in the NR1 module, while the D2 neuroreceptor was identified as a candidate provincial hub in the NR2 module (P_perm_ < 0.05; **Fig. 2F**). In contrast, the participation coefficient was used to quantify the role of neuroreceptors and transporters as connector hubs linking the NR1 and NR2 modules. The top candidate connector hubs identified were α4β2, 5-HT4, 5-HT6, CB1, and D1 (P_perm_ < 0.05; **Fig. 2G**).

Notably, the NR1 and NR2 modules were not defined by neuromodulatory families or by excitatory versus inhibitory properties. Instead, we observed distinct density distributions of these modules across the cortex (**Fig. 3**). The average density of neuroreceptors and transporters in both NR1 and NR2 was highest in the inferior prefrontal regions, insula, and temporal areas (**Fig. 3A,B**). Differences among the two modules indicated that the neuroreceptors and transporters associated with NR2 module exhibited the higher density in the motor and visual networks (P < 0.001), whereas the NR1 module exhibited the higher density in the limbic and default mode network (P < 0.001; **Fig. 3C,D**).

**Figure 3.**
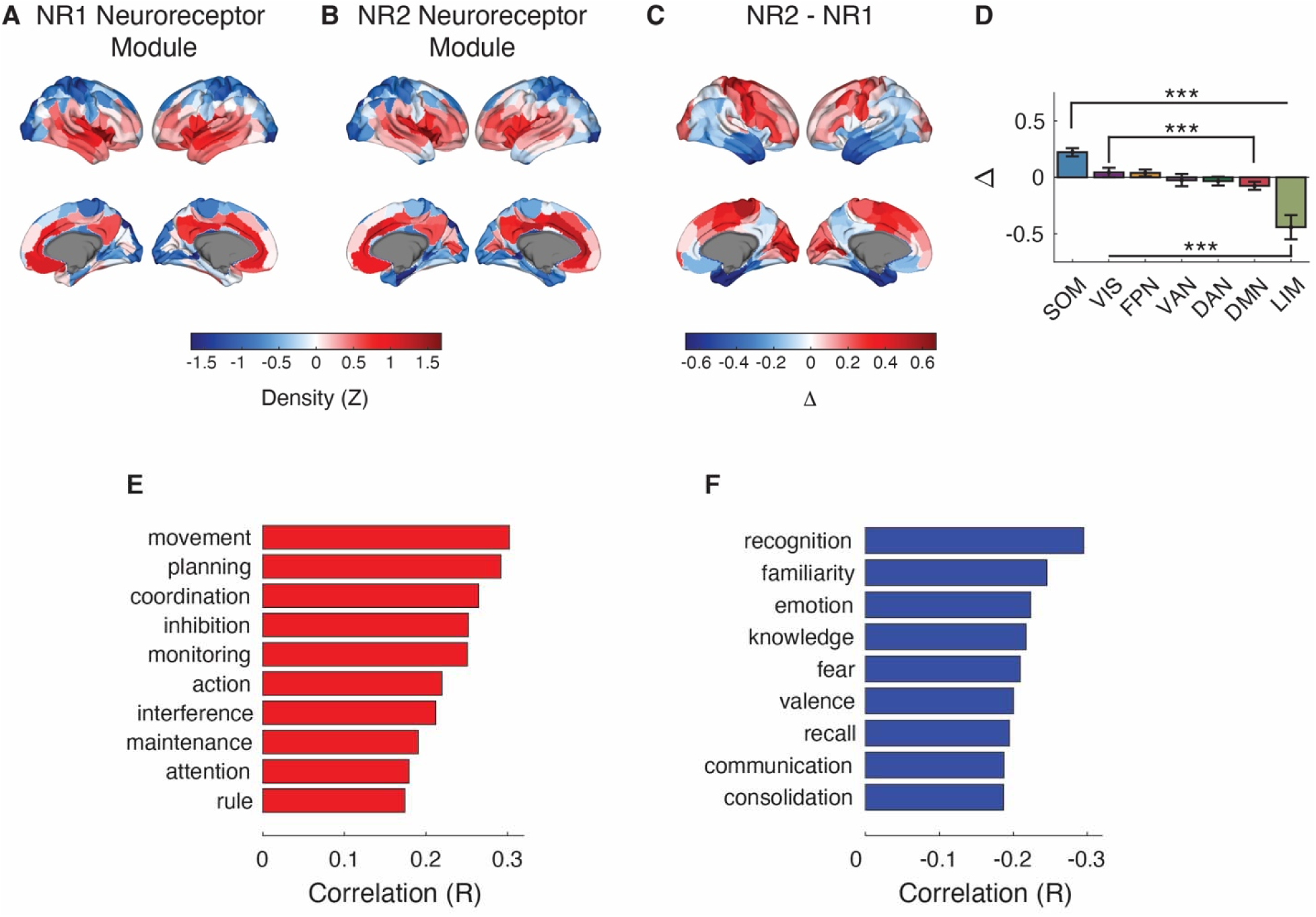
Cortical density differences between neuroreceptor modules. (A) The average density of the neuroreceptors and transporters part of the NR1 module. Module density map was estimated by averaging the z-score normalized neuroreceptor and transporter density maps part of the NR1 module. See Figure S1 for individual neuroreceptor and transporter density maps. (B) Same as panel A, but for the NR2 module. (C) Difference in the density across the cortex between the NR1 and NR2 modules. NR2-associated neuroreceptors and transporters exhibited greater density in the frontal, motor, and visual cortices, while the NR1 module was more prominent in the orbitofrontal, parietal and temporal regions. (D) Density differences between NR1 and NR2 modules among large-scale brain networks. Statistical significance was based on independent samples t-test and Bonferroni corrected for multiple comparisons. (E,F) Pearson correlation between spatial density differences, panel C, and 100 cognitive terms in Neurosynth. Only terms with significant (P < 0.05 FDR corrected) correlations are shown. FPN, Frontal Parietal Network; DMN, Default Mode Network; DAN, Dorsal Attention Network; LIM, Limbic Network; VAN, Ventral Attention Network; SOM, Somatomotor Network; VIS, Visual Network. *** P < 0.001.

These spatial differences in density between modules were associated with different cognitive processes. For the NR2 module, the cognitive processes with large positive association were “movement”, “planning” and “coordination” (P < 0.05, FDR corrected; **Fig. 3E**). On the other hand, large negative associations were observed with “recognition”, “familiarity” and “emotion” (P< 0.05, FDR corrected; **Fig. 3F**).

Lastly, the neuroreceptor drive network retained the same organization (R = 0.93) even after accounting for spatial similarity (**Fig. S3**). Additionally, the neuroreceptor drive network was consistent (R = 0.96) after regularizing the β-values using ridge regression to control for collinearity (**Fig. S4**). Together, these findings indicate that coordinated interactions among diverse neuromodulators, neuroreceptors, and transporters contribute to the shaping of emergent patterns in brain activity.

### Demonstrating consistency with known pharmacological and neurobiological associations

Merely demonstrating that the BOLD signal can be reconstructed from neuroreceptor and transporter density maps is insufficient to establish that the underlying relationship genuinely reflects neurobiological associations. This limitation highlights the necessity of providing confidence that the analysis reflects these associations. In humans, linking alterations in brain activity induced by neuropharmacological agents to known pharmacodynamics could provide such confidence. The critical aspect involves determining whether the analysis can successfully identify the neuroreceptors and transporters known to mediate the effects of a given neuropharmacological agent.

Toward this end, we utilized two datasets in which participants received LSD and Modafinil. In the first dataset, participants (N = 15) underwent 7 minutes of resting-state fMRI after receiving a Placebo and LSD^30^. In the second dataset, participants (N = 13) underwent 5 minutes of resting-state fMRI scanning before and after receiving Modafinil^31^. Moreover, understanding the mechanism of action of these drugs is important because they have been shown to exhibit beneficial effects for the treatment of neurological and neuropsychiatric disorders^32,33^.

Focusing on the LSD dataset, the BOLD signal and the functional connectivity were reconstructed in the same manner as was done in the HCP dataset. For the Placebo, the BOLD signal was reconstructed significantly greater than both null models (R_BOLD_ = 0.68 ± 0.02; P_spatial_ < 0.001; P_spin_ < 0.001) as well as the functional connectivity (R_FC_ = 0.64; P_spatial_ < 0.001; P_spin_ < 0.001) (**Fig. S5A-C**). Similar results were obtained after the administration of LSD for the BOLD signal (R_BOLD_ = 0.66 ± 0.02) and functional connectivity (R_FC_ = 0.63; P_spatial_ < 0.001; P_spin_ < 0.001; **Fig. S5D-F**). However, the model reconstructed the temporal BOLD signal significantly better after administration of the Placebo compared to LSD (t = 2.82, P = 0.01) but the differences were minor (**Fig. S5G**). Moreover, the neuroreceptor-based framework reconstructed the empirical changes in functional connectivity between the LSD and the Placebo conditions estimated using edgewise paired sample t-test significantly greater than both null models (R_LSD-Placebo_ = 0.38; P_spatial_ < 0.001; P_spin_ < 0.001; **Fig. S5H,I**).

The critical element in the analysis is to determine the neuroreceptor(s) and transporter(s) altering the neuroreceptor drive network and correspondingly the brain activity from the state before (*D_pre_*) to the state after (*D_post_*) administration of a neuropharmacological agent (**Fig. 4A**). The neuroreceptors and transporters inducing these changes are reflected in the modulation network. The modulation network is estimated from the neuroreceptor drive networks *D_pre_* and *D_post_*, which corresponds to the state before and after administration of LSD (**Fig. 4B,C**).

**Figure 4.**
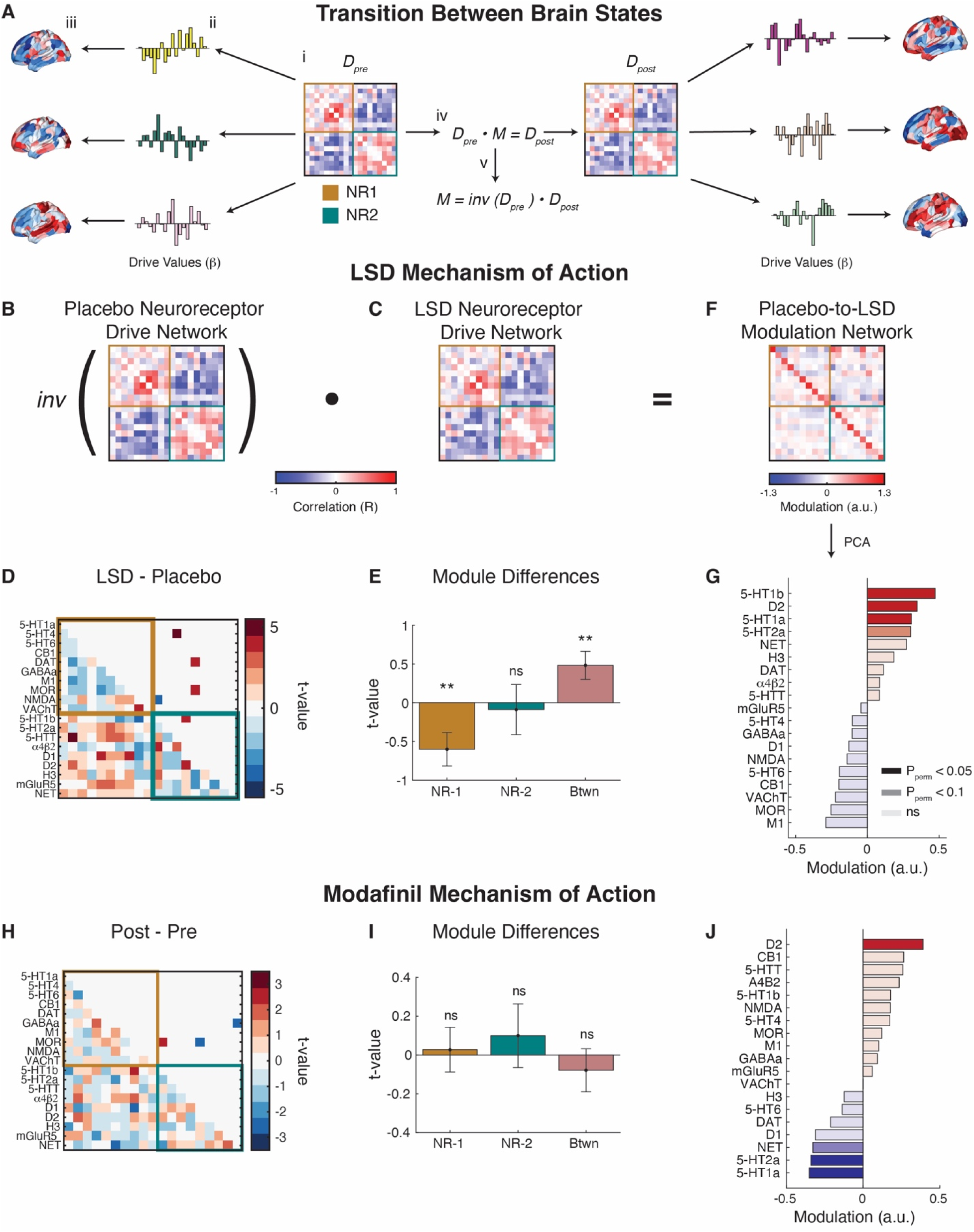
Neuroreceptor-based framework identifies the neuroreceptors and transporters mediating the effects of LSD and Modafinil. (A) Schematic representation of transition between brain states. (i) The neuroreceptor drive network, *D_pre_*, (ii) dictates the extent to which individual neurochemicals acting through their receptors contribute to (iii) brain activity. (iv) The transition between brain states, such as before to after administration of a neuropharmacological agent is mediated by a modulation network, *M*, that shifts the drive network from *D_pre_* to *D_post_.* (v) *M* can be determined by taking the inner product between the inverse of *D_pre_* and *D_post_*. Critically, all diagonal elements in the neuroreceptor drive networks are set to zero. Panel A is only for illustrative purposes and formal analyses are shown in panels B-J. (B-G) Mechanism of action of LSD. (B) Group-level neuroreceptor drive network after placebo administration, estimated from the drive values from each of the 19 neuroreceptor and transporter using Pearson correlation. Group-level neuroreceptor drive network is estimated by averaging the neuroreceptor drive network across participants. The network is organized according to the module organization identified in the HCP dataset. (C) Same as panel B, but after administration of LSD. (D) Differences between the LSD and placebo drive networks, assessed using paired samples t-tests. Correlation values were Fisher z-transformed prior to statistical testing. The top half of the network shows statistically significant changes (P < 0.05, FDR corrected), while the bottom half illustrates the magnitude of all pairwise changes. (E) Changes in drive network connectivity within and between (Btwn) modules, evaluated using a one-sample t-test to determine significant deviations from zero. (F) Group-level modulation network driving the shifts in brain activity from the placebo to the LSD-induced brain state. Modulation network is estimated by taking the inner product between the inverse of Placebo (panel B) and LSD Neuroreceptor drive network (panel C). (G) The first principal component of the modulation network to determine the extent of modulation induced by each neuroreceptor and transporter in driving the transition from the placebo to LSD state. (H-J) Mechanism of action of Modafinil. Same as panel D, E, and G, but for Modafinil. **P < 0.01; *P < 0.05.

The neuroreceptor drive network exhibited only a small number of significant pairwise changes in response to LSD (P_FDR_ < 0.05; **Fig. 4D**). These alterations were part of broader shifts within and between modules indicating significant decreases in coordination among the neuroreceptors part of the NR1 module (t_ave_ = -0.60 ± 1.45; P = 0.01) and a shift toward enhanced coordination between modules (t_ave_ = 0.48 ± 1.71; P = 0.01; **Fig. 4E**).

As can be observed in Figure 4F, the modulatory network indicated that LSD exerted a complex set of changes affecting most relationships across the neuroreceptor drive network. To determine the neuroreceptors primarily responsible for the modulations, Principal Component Analysis (PCA) was used to identify the dominant constituents underlying modulation network. The first principal component of the modulatory network accounted for 35.89% (P_perm_ < 0.001) of the variance and all other components were not significant (P_perm_ > 0.05; **Fig. S6A**). The neuroreceptor primarily responsible for driving LSD induced changes in brain activity was 5-HT1b (P_perm_ = 7.2 x 10^-4^), followed by D2 (P_perm_ = 0.01), 5-HT1a (P_perm_ = 0.03) and 5-HT2a (P_perm_ = 0.07; **Fig. 4G**). These neuroreceptors are known to bind LSD, with 5-HT1b in particular having the highest affinity for LSD^34^.

Extending the analysis to Modafinil, the model reconstructed the BOLD signal and functional connectivity *pre* (R_BOLD_ = 0.59 ± 0.03; R_FC_ = 0.62; P_spatial_ < 0.001; P_spin_ < 0.001) and *post* Modafinil (R_BOLD_ = 0.60 ± 0.03; R_FC_ = 0.66; P_spatial_ < 0.001; P_spin_ < 0.001; **Fig. S7A-F**).

Additionally, we tested if there were statistically significant differences in the reconstruction of the BOLD signal before and after Modafinil. The analysis reconstructed equally well the brain activity *pre* and *post* Modafinil (t = -1.34, P = 0.20) and the reconstructed functional connectivity reflected the observed differences (R_Post-Pre_ = 0.37; P_spatial_ < 0.001; P_spin_ < 0.001; **Fig. S7G-I**). Further, significant pairwise differences in the neuroreceptor drive network were found (P < 0.05; **Fig. 4H**; **Fig S7J,K**), but no significant differences were observed within and between the NR1 and NR2 modules (P > 0.05; **Fig. 4I**). Both the significant and non-significant pairwise differences deviate in direction within and between modules. However, these fluctuations result in an average within and between module differences not significantly deviating from zero.

The modulation network indicated that Modafinil mediated effects were driven by a subset of neuroreceptors and transporters (**Fig. S7L**). The first PCA component which accounted for 99.02% (P_perm_ < 0.001; **Fig. S6B)** of the variance in the modulation network indicated that Modafinil mediated effects were primarily driven by D2 (P_perm_ = 0.02), followed by 5-HT1a (P_perm_ = 0.03), 5-HT2b (P_perm_ = 0.05), and NET (P_perm_ = 0.07; **Fig. 4J**). The exact mechanism of action of Modafinil is not known, but these results align with prior work showing that Modafinil alters dopamine and serotonin levels in the brain^35–38^. Take together, these results provides confidence that the analysis reflects neurobiological associations.

### Neuroreceptors associated with altered BOLD signal in neuropsychiatric illness

We then investigated whether this framework could be extended to uncovered associations between neuroreceptors, transporters, and altered brain activity in neuropsychiatric disorders. Linking neuropsychiatric illnesses to specific neuroreceptors, transporters or neuromodulators is critical since it could improve diagnosis and treatment. For this analysis, we used the UCLA Consortium for Neuropsychiatric Phenomics (UCLA5c) dataset, which includes individuals diagnosed with schizophrenia (N = 45), bipolar disorder (N = 36), and attention deficit/hyperactivity disorder (ADHD; N = 39), along with a healthy control group (N = 118)^39^.

To identify the neuroreceptors and transporters associated with schizophrenia, bipolar disorder, and ADHD, we applied the same framework used to determine the mechanism of action of LSD and Modafinil. We reconstructed the temporal BOLD signal for each participant using the 19 neuroreceptor and transporter density maps. The reconstruction accuracy of the healthy control group (R_BOLD_ = 0.55 ± 0.01; mean ± SEM) was not significantly different from the schizophrenia (R_BOLD_ = 0.55 ± 0.01; P_Diff_SCZ_to_HC_ = 0.80), bipolar disorder (R_BOLD_ = 0.55 ± 0.02; P_Diff_BD_to_HC_ = 0.08), and ADHD groups (R_BOLD_ = 0.55 ± 0.01; P_Diff_ADHD_to_HC_ = 0.43; **Fig. S8**). Additionally, the neuroreceptor-based framework reconstructed differences in functional connectivity between neuropsychiatric groups and healthy controls significantly greater than null models for schizophrenia (R_SCZ-HC_FC_ = 0.56; P_spatial_ < 0.001; P_spin_ < 0.001), bipolar disorder (R_BD-HC_FC_ = 0.45; P_spatial_ < 0.001; P_spin_ < 0.001), and ADHD (R_ADHD-HC_FC_ = 0.35; P_spatial_ < 0.001; P_spin_ < 0.001; **Fig. S9**). In the schizophrenia group, the neuroreceptor drive network exhibited a limited number of significant pairwise changes compared to healthy controls (P < 0.05) and these changes were part of a broader set of alterations within and between neuroreceptor drive network modules (P < 0.05; **Fig. S10A**).

To determine the neuroreceptors and transporters driving these changes, we estimated the modulation network required to transition from the healthy to the schizophrenic brain state in the same manner as the analysis on LSD and Modafinil. The modulation network exhibited a complex pattern, with the first principal component accounting for 67.65% of the variance (P_perm_ < 0.001); all other components were not significant (P_perm_ > 0.05; **Fig. S10B**). The modulation analysis identified norepinephrine transporter (NET) as the primary driver (P_perm_ = 0.02; **Fig. 5A**). Other neuroreceptors mediating the effects included serotonin receptors 5-HT1a (P_perm_ = 0.03) and 5-HT4 (P_perm_ = 0.09), and additional associations with NMDA (P_perm_ = 0.02), α4β2 (P_perm_ = 0.07), and GABAa (P_perm_ = 0.09) were observed.

**Figure 5.**
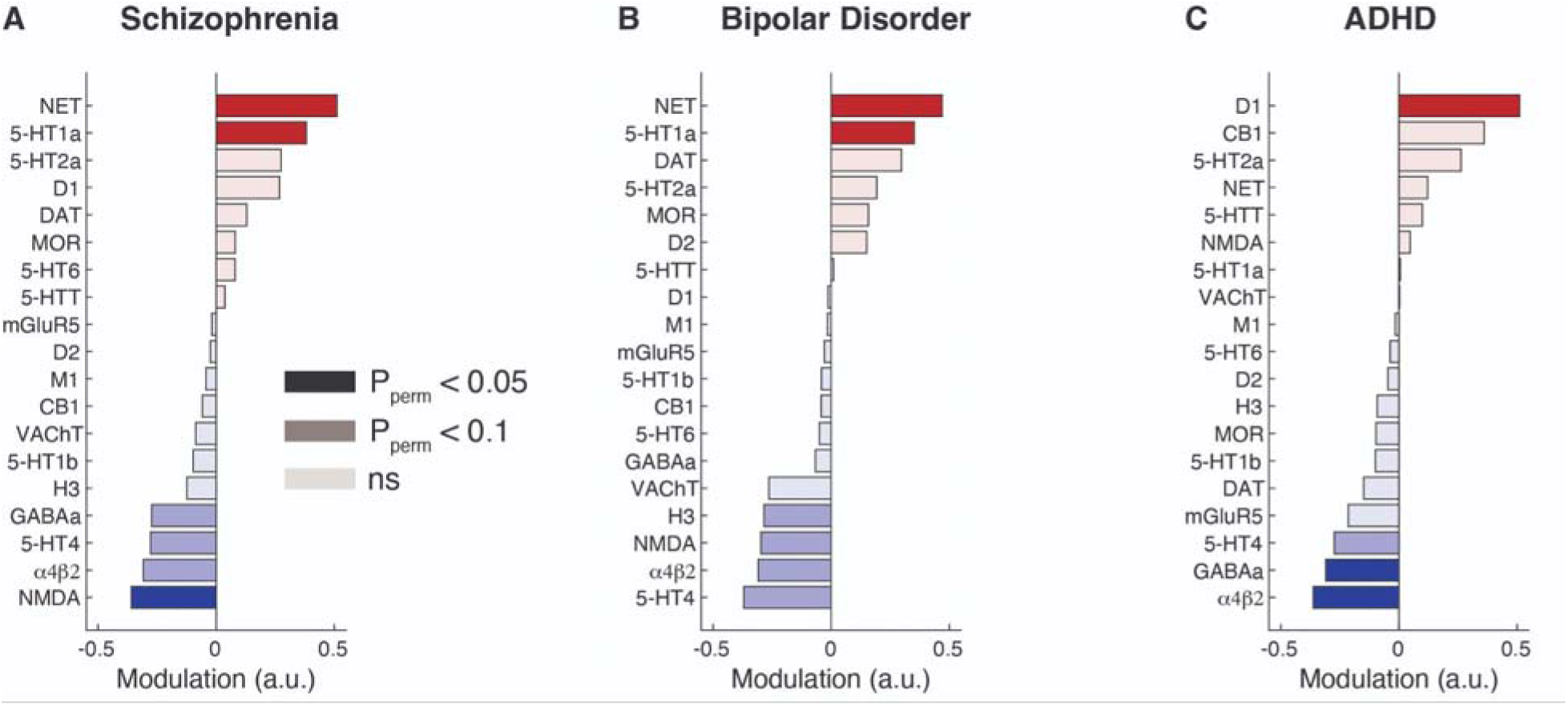
Neuroreceptors and transporters mediating the alterations in brain activity across neuropsychiatric disorders. (A) Neuroreceptors and transporters mediating the changes in brain activity from healthy controls (HC) to schizophrenia (SCZ). (B,C) Same as panel A, but for bipolar disorder (BD) and ADHD, respectively. Statistical significance was determined based on a permutation test and P_perm_ < 0.05 were considered significant. ns, not significant.

For bipolar disorder and ADHD, the analysis similarly identified a small set of pairwise changes within and between neuroreceptor network modules (P < 0.05; **Fig. S10C-F**). In bipolar disorder, NET was the primary driver similar to schizophrenia (P_perm_ = 0.04; **Fig. 5B**). Overall, neuroreceptor mediating the effect of bipolar disorder included 5-HT1a (P_perm_ = 0.01), 5-HT4 (P_perm_ = 0.06), α4β2 (P_perm_ = 0.07), NMDA (P_perm_ = 0.08), and H3 histamine (P_perm_ = 0.09). In ADHD, changes were mediated by the dopamine, D1 (P_perm_ = 0.01), and acetylcholine, α4β2 (P_perm_ = 0.01; **Fig. 5C**). The analysis also implicated GABAa (P_perm_ = 0.04) and the 5-HT4 serotonin neuroreceptor (P_perm_ = 0.07). Taken together, these findings underscore the complexity of neuropsychiatric illness.

### Consistent neuroreceptor drive networks across datasets

Further, we assessed the similarity of neuroreceptor drive networks across datasets. Strong similarity between these independent datasets would support the robustness of the results and framework. To quantify the extent of similarity, we compared the neuroreceptor drive networks derived from the LSD, modafinil, and UCLA5c datasets to the network estimated from the HCP dataset. Neuroreceptor drive networks from each of the other datasets showed strong similarity with the HCP-derived network (R_ave_ = 0.92; **Fig. S11**). These results underscores the stable dynamical interplay between brain activity and the underlying neuromodulator, neuroreceptors and transporters.

### Reconstructing spontaneous brain activity: neuroreceptors vs. structural and geometric modes

Lastly, previous work has shown that spontaneous brain activity is shaped and constrained by anatomical factors – structural connectivity (SC)^40,41^ and cortical folding geometry (GC)^42^. Models based on SC and GC suggest that spontaneous brain activity emerges from a weighted linear summation of modes (also called harmonics) of the SC or GC. However, it is an open question as to which model – neuroreceptor, structural or geometric – can more parsimoniously reconstruct spontaneous brain activity.

To compare models, we utilized the BOLD signal from the HCP dataset due to its extensive scanning duration. PCA was applied to the concatenated BOLD signal to identify the dominant spatial patterns of activity. The analysis utilized components C1 through C10 which account for 40.85% of variance in the BOLD signal. We opted to do the analysis on the components of the BOLD signal because this balanced the number of comparisons (i.e. components) and interpretability. Specifically, using the components C1-to-C10, we were able to account for 40.85% of BOLD signal while preserving interpretability since there are only ten spatial maps. On the other hand, using the timeseries of the BOLD signal would have increased the temporal resolution but it would have come at the cost and challenge of interpreting the spatial patterning which the anatomical modes are more parsimoniously able to reconstruct the BOLD signal. Additionally, since PCA is a linear decomposition of the spatial temporal BOLD signal, it preserves the global structure of the data. Specifically, the linear mapping preserves the underlying distances in lower dimensional space compared to non-linear methods such Uniform Manifold Approximation (UMAP) or t-SNE in which the distances in the lower dimensional space are distorted.

Specifically, we determined the number of SC- and GC-modes needed to match the accuracy obtained with the neuroreceptor maps. SC or GC based reconstructions were deemed to be more parsimonious if fewer modes were required to achieve the same accuracy as the neuroreceptor maps. The model comparison revealed that reconstructing the BOLD signal using the neuroreceptor maps was more parsimonious for 8 out of the 10 BOLD components (**Fig. S12, S13**). These results suggest that the neuroreceptor-based framework may provide a more direct and biologically meaningful basis for modeling brain activity compared to anatomical-based approaches.

## Discussion

Neurotransmitters are the primary substrate by which neurons communicate^43^. Like most aspects of the human brain, no individual neuromodulator works alone to generate brain activity; instead brain activity is generated from coordination among multiple neuromodulators acting through their corresponding neuroreceptors and transporters^27^. Here, we present a neuroreceptor-based framework to identify the neuroreceptors and transporters working in congruence to drive spontaneous brain activity. Across four datasets, the neuroreceptor-based framework identified consistent associations between neuroreceptors and transporter density maps and fluctuations in the BOLD signal.

Spontaneous brain activity reflected a coordinated interplay among a diverse set of excitatory and inhibitory neuroreceptors and transporters, highlighting the complex neuromodulatory landscape of the cortex. This finding raises the hypothesis that the balance between excitation and inhibition is managed by two distinct modules – NR1 and NR2 – reflecting visual and somatomotor networks or higher-order associative functions, respectively. The differing density distributions between modules across cortical regions suggests that brain activity and function is locally shaped by specific neuromodulatory profiles. For instance, the higher density of NR1 module in orbitofrontal, parietal and temporal areas may facilitate the integration of sensory and associative information^44^, whereas the NR2 module’s prominence in frontal, motor, and visual regions may align with the demand for rapid modulation of motor responses and visual processing^45,46^. This spatial organization suggests that the brain dynamically allocates neuromodulatory resources in a region-specific manner which may enhance its capacity to adapt to diverse cognitive and behavioral demands. Moreover, the modules may serve as a safeguard against excessive excitation synchronization across cortical regions^47,48^.

Why has the brain gone to such a great extent to maintain a rich tapestry of neuromodulators, neuroreceptors and transporters to facilitate communication? One possible reason could be that utilizing multiple neuromodulators, neuroreceptors and transporters could fine tune the excitation-inhibition balance, enabling neurons to maintain responsiveness across a broad spectrum of inputs and to adapt to varying inputs without changing their fundamental properties^49,50^. Furthermore, the interplay between different neuromodulators, neuroreceptors, and transporters may serve an important role in network-level flexibility. We speculate that the ability to respond to diverse neuromodulators enables a brain region to be part of multiple subnetworks depending on the context. This versatility may facilitate efficient communication across the brain and support diverse cognitive functions without necessitating dedicated, fixed circuitry for every task^51^. Such a flexible integration among multiple subnetworks could allow the brain to maximize computational power while minimizing the metabolic costs^52^.

The analysis also provided insight into the mechanism of action of an administered neuropharmacological agent. Determining the mechanism of action has been challenging because neuropharmacological agents have direct and indirect effects on the activation propensity of neuroreceptors and neurons^53^. The analysis identified the neuroreceptors that bind LSD and neuroreceptors thought to mediate the effects of Modafinil, providing support for known pharmacological and neurobiological associations. The analysis identified 5-HT1b as having high binding affinity for LSD in line with previous work^34^ and suggesting the need for additional investigation of non-5HT2a-mediated mechanisms^54^. However, there is a complex relationship between binding affinity and psychopharmacology. For instance, 5-HT2a is thought to be the primary functional mediator of psychedelic effects of LSD suggesting that binding affinity may not directly translate to functional significance^55,56^.

We extended our framework to investigate neuropsychiatric disorders—schizophrenia, bipolar disorder, and ADHD—to identify the neuroreceptors and transporters associated with alterations in brain activity for each condition. Specifically, in line with prior work, in schizophrenia the analysis identified significant associations with NET^57^, 5-HT1a^58^, NMDA^59–61^ and trending associations with α4β2^62,63^, 5-HT4 and GABAa^64^. Interestingly, our results did not identify a significant association with dopamine neuroreceptors and transporters aligning more closely with alternative hypotheses than the current dominant dopamine hypothesis in schizophrenia^65^. The neuroreceptors associated with bipolar disorder were similar to the ones associated with schizophrenia indicative of their close psychopathology^66^. In the ADHD group, consistent with previous work, alteration in brain activity were associated with the D1^67–69^, α4β2^70–73^, GABAa^74,75^, and 5-HT4^76^ neuroreceptors. However, unlike prior work, we did not find an association with norepinephrine system^77^. (Note see Supplemental for extended discussion).

Together, the results suggest that these disorders may involve dysregulation across multiple neuromodulators and/or neuroreceptor and transporter systems rather than a single system. This dysfunction affecting multiple neuroreceptor systems could arise from an initial impairment in one system, with compensatory adjustments in others to maintain homeostasis^78^ or it could be indicative of a common underlying factor disrupting multiple systems^79^. Alternatively, the observed dysfunction may reflect underlying abnormalities in the neurons responsible for releasing neurotransmitters such as norepinephrine or dopamine, with downstream effects manifesting in the associated neuroreceptors and transporters. The pattern of dysfunction spanning multiple systems highlights the complexity of neuropsychiatric conditions, underscoring the need for integrative approaches that consider the interplay among diverse neuromodulators. Toward this end, multimodal imaging can deepen our understanding of the coordinated roles of neuroreceptors and transporters in maintaining healthy brain function^80^ and how their dysregulation can lead to pathological states^81,82^.

Moreover, the framework could be utilized to provide insight into the mechanism of action of other neuropharmacological agents and narrow the search for therapeutic treatments for substance abuse^83,84^ and neuropsychiatric illness^85,86^. For instance, the findings in the psychiatric group could be used to identify compounds and/or optimal combinations of existing medications to improve the efficacy. Overall, caution is warranted when interpreting these results given the small sample size and that there is no ground truth in the molecular factors underlying these disorders. The current work has several limitations. First, causal relationship cannot be drawn from the analysis because the analysis is correlational. Investigating the spatial pattern of a neuroreceptor is insufficient to draw conclusions on the mechanism of action of neuropharmacological agents or neuropsychiatric illness. Specifically, the mechanism of action can be affected by multiple factors such as affinity, expression levels on different neuronal layers or downstream effects. Moreover, the assumption that a certain neurotransmitter affects the BOLD signal in the exact same region may not hold considering the complex projection of modulatory neurotransmitter systems. Future studies can utilize simultaneous PET/fMRI imaging to establish causal relationships. Second, despite the neuroreceptor frameworks’ ability to reconstruct the BOLD signal greater than both null models, this still leaves a large portion of spatial variance unexplained. This could in part be due to the small number of neuroreceptors used in the analysis. On the one hand the 19 neuroreceptor and transporter density maps used in the analysis are the most comprehensive maps to date, on the other hand these represents only a fraction of the total number in the brain. Therefore, future studies can test if the inclusion of additional neuroreceptors increases the variance explained in the BOLD signal. Third, the neuroreceptor and transporter maps have different spatial resolutions, smoothing levels, extent of autocorrelation, and scanner of origin which could bias their correlation with BOLD signal. Further, the spatial and spin null models used in the analysis may account for some aspect of these differences. Specifically, the spin null model may account for the autocorrelation in the neuroreceptor maps, nonetheless, it is an open question to the extent these null models account for these underlying factors. Future studies could account for this by using neuroreceptor maps from a single source or harmonizing the pre-processing of PET images to mitigate such concerns. Lastly, the analysis focused on cortical regions which limits the biological interpretation, especially given that many key neuroreceptor systems have high density in subcortical structures. These subcortical regions may be critical in mediating alteration in brain activity due to psychiatric illness, medication or psychedelics^87^. This could explain the lack of association between neurotransmitter systems like dopamine in schizophrenia. Therefore, future studies can incorporate neuroreceptor maps from subcortical regions to provide a holistic relationship between molecular factors, brain activity and neuropsychiatric illness.

In summary, identifying the neuromodulators that drive brain activity is crucial due to their central roles in communication, cognition, and pathology. Here, we demonstrated that using cortical density maps of neuroreceptors and transporters from PET imaging can reveal the underlying neuroreceptors and transporters underlying spontaneous brain activity. In the process, the framework revealed a complex network of neuroreceptors, and transporters associated with brain activity. Further, this neuroreceptor-based framework can improve our ability to identify neuromodulators driving brain activity and opens new avenues for exploring dynamical brain activity, identifying neuropharmacological mechanisms of action and markers of neuropsychiatric disorders. Future studies can build on these findings to unravel the complex mechanisms that underlie cognition, and in generating targeted diagnostics and therapies.

## Methods

### Neuroreceptor Density Maps

The analysis used data originally published in Hansen et al., 2022^27^. In brief, PET tracer imaging was used to map the density for 19 neuroreceptors and transporters. The neuroreceptors include 9 neurotransmitter systems: dopamine (D1, D2, DAT), norepinephrine (NET), serotonin (5-HT1A, 5-HT1B, 5-HT2A, 5-HT4, 5-HT6, 5-HTT), acetylcholine (α4β2, M1, VAChT), glutamate (mGluR5, NMDA), γ-aminobutyric acid (GABAa), histamine (H3), cannabinoid (CB1), and μ-opioid neuroreceptor (MOR). PET images were acquired only from healthy participants (*n* = 1,238; 718 males and 520 females). The images are used to estimate the binding potential and tracer distribution volume which are referred to as receptor densities. PET based estimation methods provide a direct measure of the neuroreceptor and transporter concentration across the cortex than estimates based on transcriptomics^74^. PET images were all registered to the MNI-ICBM 152 non-linear 2009 template and parcellated into the 200 regions according to the Schaefer atlas^28^. Finally, each map corresponding to individual neuroreceptors and transporters were z-score normalized. Moreover, the neuroreceptor maps, which are z-scored before being related to BOLD activity, are treated as maps of relative spatial distribution of neuroreceptor/transporter density. Note that the ranking across regions is the same, whether an absolute estimate (e.g. distribution volume) or relative estimate (e.g. distribution volume ratio). What may change the ranking of regions is the presence of flow effects or non-specific binding (e.g., when using a semiquantitative SUVR instead of specific volume or binding potential).

### Functional MRI Data

#### HCP Resting-State fMRI Dataset

We used pre-processed resting-state functional MRI (fMRI) data from 48 healthy human participants from the Human Connectome Project (HCP)^29^. We downloaded 50 participants, but two were missing data. In brief, participants underwent four sessions of 15-min resting-state scanning sessions. fMRI volumes were recorded using a customized 3T Siemens Connectome Skyra scanner with an EPI sequence (TR = 0.72 s, TE = 33.1 ms, 72 slices, 2.0 mm isotropic, field of view (FOV) = 208 × 180 mm). In addition to the pre-processing steps part of the HCP pipeline, we additionally regressed out the global signal, white matter, and cerebral spinal fluid. The global signal was removed in order to reconstruct the aspect of the BOLD signal that more closely reflets neuronal activity since the global signal is associated with motion and/or physiological noise^88^. The application global signal regression (GSR) to the fMRI data is an ongoing debate and different associations might be observed with the receptor templates if GSR is not applied^89^. The data was mapped on Schaefer atlas to derive brain activity for 200 regions of interest because voxel-wise estimates can be unstable and noisy. Group-level functional connectivity (FC) was estimated using the Pearson correlation on the pooled data.

#### Lysergic Acid Diethylamide (LSD) Resting-State fMRI Dataset

The analysis was based on data detailed in Carhart-Harris et. al., 2016^30^. In brief, the analysis utilized pre-processed fMRI data from 15 participants that underwent 7 minutes of resting-state fMRI after a placebo and LSD. fMRI data were acquired using a gradient echo planar imaging sequence, TR/TE = 2000/35ms, field-of-view = 220mm, 64 × 64 acquisition matrix, parallel acceleration factor = 2, 90° flip angle and 35 oblique axial slices and 3.4mm^3^ voxel size.

Pre-processing steps are detailed in Carhart-Harris et. al., 2016. fMRI data pre-processing involved de-spiking, slice time correction, motion correction, non-linear registration to 2mm MNI brain. Additionally, fMRI volumes with frame displacement greater than 0.4mm were removed. Finally, the functional data were spatially smoothed with a 6mm full-width half max kernel and band-pass filtering between 0.01 to 0.08 Hz. Finally, 9 nuisance regressors (6 motion-related and 3 were anatomically-related) and linear and quadratic trends were removed. As in the HCP dataset, we additionally regressed out the global signal. Subject-level functional connectivity was estimated using the Pearson correlation.

#### Modafinil Resting-State fMRI Dataset

The analysis was based on data detailed in Cera et. al., 2014^31^. In brief, the study recruited 26 young male right-handed (as assessed by the Edinburgh Handedness inventory) adults (age range: 25–35 y.o.). Participants received, in a double-blind fashion, either a single dose of Modafinil or a placebo pill identical to the drug. The present analysis focused only on 13 participants that received Modafinil.

Resting-state-fMRI BOLD data were separated in three runs lasting four minutes each followed by high resolution T1 anatomical images. BOLD functional imaging was performed with a Philips Achieva 3T Scanner (Philips Medical Systems, Best, The Netherlands), using T2*-weighted echo planar imaging (EPI) free induction decay (FID) sequences and applying the following parameters: TE 35 ms, matrix size 64×64, FOV 256 mm, in-plane voxel size 4×4 mm, flip angle 75°, slice thickness 4 mm and no gaps. 140 functional volumes consisting of 30 transaxial slices were acquired per run with a volume TR of 1671 ms. High resolution structural images were acquired at the end of the three rs-fMRI runs through a 3D MPRAGE sequence employing the following parameters: sagittal, matrix 256×256, FOV 256 mm, slice thickness 1 mm, no gaps, in-plane voxel size 1 mm×1 mm, flip angle 12°, TR = 9.7 ms and TE = 4 ms.

MRI data were preprocessed with the CONN Toolbox^90^. The preprocessed with the following steps: de-spiking, slice-timing correction, realignment, segmentation, coregistration, normalization, and spatial smoothing with 8 mm full width half maximum (FWHM). The preprocessing of the T1-weighted structural images involved skull-removal, normalization into MNI anatomical standard space, and segmentation into gray matter, white matter, and cerebral spinal fluid, soft tissues, and air and background. In functional images, all volumes with frame displacement greater than 0.3 mm were censored and followed by regression of the global signal.

#### UCLA Consortium for Neuropsychiatric Phenomics Dataset

The Consortium for Neuropsychiatric Phenomics includes imaging of healthy individuals (N=118 out of 122), individuals diagnosed with schizophrenia (N= 45 out of 50), bipolar disorder (N=35 out of 37), and attention deficit hyperactivity disorder (N= 39 out of 40)^39^. Resting-state functional MRI scans were obtained while participants kept their eyes open for 5 minutes in the scanner. Neuroimaging data were acquired on a 3T Siemens Trio scanner. T1-weighted high-resolution anatomical scans (MPRAGE) were collected with a slice thickness = 1mm, 176 slices, TR=1.9s, TE=2.26ms, matrix=256 x 256, FOV=250mm. Resting-state MRI data were collected with a T2*-weighted EPI sequence with slice thickness = 4mm, 34 slices, TR=2s, TE=30ms, flip angle=90°, matrix=64 × 64, FOV=192mm.

The analysis was based on minimally preprocessed data using FMRIPREP version 0.4.4 (http://fmriprep.readthedocs.io). T1-weighted volume was corrected for bias field using ANTs N4BiasFieldCorrection v2.1.0 skullstripped and coregistered ICBM 152 Nonlinear Asymmetrical template. Resting-state data were motion corrected using MCFLIRT v5.0.9. Functional data was skullstripped and coregistered to the corresponding T1-weighted volume using boundary-based registration. Motion correcting transformations, transformation to T1-weighted space and MNI template warp were applied in a single step using antsApplyTransformations. Framewise displacement and dvars were calculated using Nipype implementation. In addition to those regressors, global signal and mean white matter signal was also calculated and removed. Additionally, analysis was conducted only on participants with an average frame displacement less than 0.5 mm. In total the analysis was based on 118 out of 130 healthy controls, 45 out of 50 participants diagnosed with schizophrenia, 35 out of 49 participants diagnosed with bipolar disorder and 39 out of 43 participants diagnosed with ADHD. Moreover, in the remaining participants, all individual fMRI volumes with frame displacement greater than 0.3mm were removed. This dataset was chosen because it encompasses a diverse range of neuropsychiatric disorders while minimizing the potential measurement discrepancies that can occur when combining data from multiple sources.

### Reconstructing BOLD Time Series, Components and Functional Connectivity from Neuroreceptor Density Maps

Reconstruction of the BOLD signal was conducted on both the timeseries and principal components of the BOLD signal. The same method was used for reconstructing the BOLD time series. Specifically, each of 1,200 fMRI volumes (time points) per participant were reconstructed from a weighted linear summation of the 19 neuroreceptor and transporter density maps.

Formally, each volume of the BOLD signal is reconstructed as:

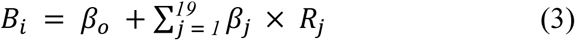

Where *B_i_* is the *i^th^* BOLD volume. Further, *R_j_* is the spatial density map associated with the *j^th^* neuroreceptor and transporter, *β_o_* is the intercept and *β_j_* is the estimated beta-values for each neuroreceptor map in the model. Model fitting was performed using *fitlm.m* in Matlab 2021b.

Reconstruction accuracy of the BOLD signal was estimated using the Pearson correlation between empirical and reconstructed components. We limited our analysis to the cortex because there are additional challenges associated with estimating the subcortical BOLD signal. For instance, subcortical BOLD signal tends to have lower signal-to-noise ratio^91^.

Linear regression beta estimates rely on the independence of the model terms. However, when terms exhibit collinearity, the least-squares estimate can be unstable. To test the extent this issue affected our results, we used ridge regression to regularize the beta coefficients. For the analysis, we compared the estimated neuroreceptor drive network without regularization and ridge regression normalized.

Additionally, we determined if the neuroreceptor-based reconstruction of the BOLD timeseries preserved the inherent functional connectivity present in resting-state fMRI. To determine the accuracy in the functional connectivity of the reconstructed BOLD signal, the empirical and reconstructed BOLD signal was concatenated across participants and Pearson’s correlation was used to estimate the functional connectivity, respectively.

The same framework was applied to the top ten BOLD spatial components in HCP dataset, LSD, Modafinil and UCLA5c datasets. For the component analysis, the BOLD data from all participants was pooled together resulting in 200 x 57,600 data points. Principal components of the BOLD signal were identified using *pca.m* in Matlab 2021b. Consistent with the terminology used by Pang et al., 2023, we refer to this process as "reconstruction" to describe the estimation of BOLD activity from the model^42^.

Pre-processing choices can bias fMRI results^92^. We found that that average reconstruction accuracy was comparatively similar across the four datasets (r = 0.5-to-0.6). Moreover, as shown in Fig. S11, the similarity in neuroreceptor drive networks derived from the LSD, Modafinil and UCLA5c datasets exhibited a strong similarity with the HCP-derived network (R_ave_ = 0.92). Together these results suggest that reconstructing the BOLD signal from neuroreceptor maps was robust to pre-processing choices.

### BOLD Reconstruction Null Models

#### Spatial Permutation Null Model

Reconstruction accuracy was compared to the spatial permutation null model. Specifically, for each of the 19 neuroreceptor and transporter density maps were spatially permuted and BOLD component and time series signal was reconstructed from the permuted maps. A total of 1000 permutations were conducted. Additionally, we estimated the functional connectivity from the reconstructed BOLD signal from the spatial null model by first concatenating the reconstructed BOLD signal estimated from the spatially permuted null model across participants and Pearson’s correlation was used to estimate the functional connectivity. A total of 1000 permutations were performed and observed accuracy values greater than the spatial null model at a P_spatial_ < 0.05 were considered statistically significant.

#### Spin Null Model

Additionally, reconstruction accuracy was compared to the spin permuted null model^93^. Spin permutation, randomized each of the 19 neuroreceptor and transporter density maps but preserves the distance dependencies between brain regions. Similar to the spatial permutation null model, BOLD component and time series signal was reconstructed from the spin permuted maps. In the same manner as the spatial permutation null model, we estimated the functional connectivity from the reconstructed BOLD signal from the spin null model. A total of 1000 permutations were performed and observed accuracy values greater than the spin null model at a P_spin_ < 0.05 were considered statistically significant.

### Neuroreceptor Drive Network Estimation and Network Properties

#### Estimating Neuroreceptor Drive Network

Neuroreceptor drive network was estimated by correlating the beta-values from the time series model. The neuroreceptor drive network was estimated from the pooled data across participants. Clustering of the neuroreceptor drive network was conducted using modularity-maximization^94^. Modularity-maximization does not require the number of clusters to be specified and the resolution of the clusters was controlled with resolution parameter, γ = 1. Modularity-maximization was implemented with the Generalized Louvain algorithm part of the Brain Connectivity Toolbox^95^. Clustering results from the Generalized Louvain algorithm can be dependent on the initial seeding. Therefore, clustering was repeated 100x with different initial seeds, and final clustering results were based on the consensus across iterations^96^. Specifically, consensus clustering identifies a single representative partition from the set of 100 iterations. This process involves the creation of a thresholded nodal association matrix which describes how often nodes were placed in the same cluster. The representative partition is then obtained by using a Generalized Louvain algorithm to identify clusters within the thresholded nodal association matrix.

#### Provincial Hubs

Provincial hubs are nodes in a network that have strong connection within a module^95^. A provincial hub score for each node was estimated using the within module degree which quantifies the strength of the connections a node exhibits with other nodes within a module. This involved first thresholding the network to remove all negative connections. Brain connectivity toolbox was used to estimate provincial hub scores^95^. Within-module z-score can me estimated as:

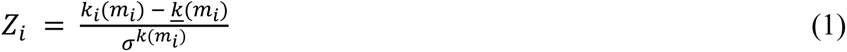

where *m_i_*is the module containing node *i*, *k_i_*(*m_i_*) is the within-module degree of *i*, and *k*(*m_i_)* and 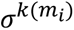 are the mean and standard deviation.

#### Connector Hubs

Connector hubs are nodes that link modules. A connector hub score was estimated for each node using participation coefficient (PC) as:

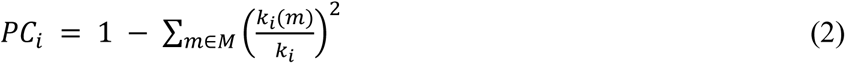

where *M* is the set of modules, and *k_i_* (*m*) is the strength of connection between *i* and all nodes in module *m*. As with the provincial hubs, the neuroreceptor network was first thresholded to remove all negative connections. Provincial hubs may act as coordinators within a module whereas connector hubs modulate the contributions between modules and brain activity. Brain Connectivity Toolbox was used to estimate participation coefficient and values were z-score normalized^95^.

#### Statistical Testing of Hub Properties

Statistical significance for both provincial and connector hub scores was based on comparison to a null model. Specifically, a degree persevering permutation was utilized to randomize the neuroreceptor network prior to estimating the provincial and connector hub scores from the null model. A total of 1000 permutations were performed and empirical hub scores that were greater than the null model at a P_perm_ < 0.05 were considered statistically significant.

#### Neurosynth Associated Cognitive Processes

Association between brain activity and cognitive processes were obtained from Neurosynth^97^. Neurosynth contains more than 2,000 terms associated with a cognitive process. We limited our analysis to the top 100 terms of interest reflecting cognitive and behavioral terms. Specifically, the brain maps representing differences density between neuroreceptor modules were correlated with spatial maps of representing the 100 cognitive terms. The analysis was performed using NIMARE^98^.

### Neuroreceptors Underlying LSD, Modafinil and Neuropsychiatric Illness Induced Changes

To identify the neuroreceptors and transporters contributing to changes in brain activity following LSD, Modafinil, and in underlying neuropsychiatric conditions, we determined the neuroreceptors and transporters mediating these effects. The key analytical step involved identifying a modulatory network, *M*, that transforms the neuroreceptor drive network from state *p* to state *q*. The relationship can be expressed mathematically as:

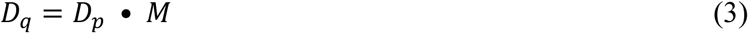

Critically, all diagonal elements in *D* are set to zero ensuring that the neuroreceptor and transporter does not activate itself in the framework. *M* can be determined by solving:

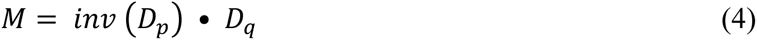

The neuroreceptors driving the modulations can be identified by decomposing *M* using PCA. Moreover, the same analysis was used to gain insight into the neuroreceptor(s) and transporter(s) underlying Modafinil and neuropsychiatric illness. Conceptually, the PC1 of the modulation network can be thought as reflecting the neuroreceptors driving the alterations in the BOLD signal due to LSD, Modafinil or psychiatric illness.

The modulatory network, *M*, can fit any difference in the neuroreceptor drive networks even if the underlying differences are simply noise. Statistical significance was based on randomizing *M*, generating a random modulatory network, *M_rand_* in a degree preserving manner. Randomizing the empirically derived *M* ensures that inherent properties such as the mean and standard deviation of *M* are preserved in *M_rand_*. Once *M_rand_* is estimated, it is decomposed using PCA and the variance explained by each component is compared to *M.* The modulation values associated with each neuroreceptor, and transporter are compared. A total of 1000 permutations are performed and the principal component and neuroreceptors and transporters in the empirically derived *M* that deviate from *M_rand_*at probability of P_perm_ < 0.05 are considered to be statistically significant.

### Model Comparison: Structural and Cortical Geometric Eigenmodes

#### Structural Connectivity and Structural Eigenmodes

Preprocessed diffusion weighted images for each of the 48 participants part of the HCP were used to construct structural connectomes. Specifically, fiber tracking was done using DSI Studio with a modified FACT algorithm^99^. Data were reconstructed using generalized q-sampling imaging (GQI) in MNI space. Fiber tracking was performed until 250,000 streamlines were reconstructed with default parameters. For each participant, an undirected weighted structural connectivity matrix, A, was constructed from the connection strength based on the number of streamlines connecting two regions. Connectivity matrix was normalized by dividing the number of streamlines (T) between region *i* and *j*, by the combined volumes (*v*) of region *i* and *j*:

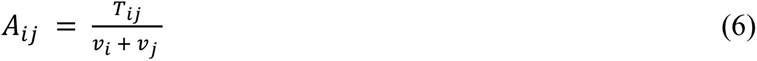

Group structural connectivity was estimated by averaging all the individual structural connectivity networks together.

In accordance with Naze *et al.*, 2021, structural eigenmodes were estimated from the normalized graph Laplacian of the structural network as:

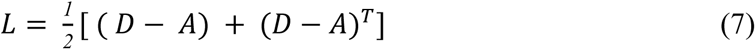

where *A* is the adjacency matrix of the group-level structural connectome and *D* is the degree matrix^100^. The graph Laplacian *L* is then decomposed into a finite number of eigenvalues λ_*k*_ and eigenvectors, ψ_*k*_:

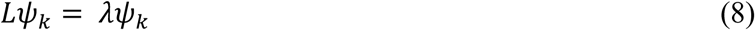

#### Cortical Geometric Eigenmodes

Cortical eigenmodes capture how the anatomy constrains brain dynamics. The analysis relies on cortical modes derived in Pang et. al., 2023^42^. In brief, cortical eigenmodes are estimated from the midthickness surface mesh part of the FreeSurfer’s fsaverage population-averaged template. Local vertex-to-vertex spatial relations and curvature are modeled using the Laplace–Beltrami operator from the surface mesh, and from which the eigenmodes are estimated. Each eigenmode comprises spatial patterns with a specific wavelength.

#### Comparing Reconstruction Models

To compare the reconstruction models, we use the first 10 principal components of the BOLD signal. Specifically, each component is reconstructed using structural and geometric eigenmodes by sequentially adding modes to the model until the reconstruction accuracy of the models equals the accuracy obtained with the neuroreceptor-based model. Structural- or cortical geometric-based reconstructions were deemed to be more parsimonious if fewer modes were required to achieve the save accuracy as the neuroreceptor-based framework.

We can only compare explained variance for a given set of input variables instead of the maximally explained variance by a model because the number of neuroreceptor maps is limited by the number of available PET imaging data on neuroreceptors; in this case 19. Whereas the modes are limited by the dimensionality of the data. For instance, 200 in case of structural connectivity. Therefore, adding more modes to a model could inherently increase the explainable variance of the BOLD signal. Nonetheless, there is still value in conducting the comparison since the preferred model would be the one that can explain the same amount of variance with the fewest parameters.

### Statistics and Reproducibility

All statistical testing was conducted using Matlab 2021b unless otherwise stated. Linear modeling was conducted using *fitlm.m* in Matlab. All neuroreceptor drive network were estimated using Pearson correlations of the β-coefficients estimated using *fitlm.m*. Spatial permutations and spin permutated maps were generated in Matlab (see section on BOLD Reconstruction Null Models for details). Differences were assessed using Student’s *t*-test and FDR corrected for multiple comparison. Detailed statistical information, including exact *n* values and test types, is provided in the respective figure legends. Statistical parameters, including mean and standard error of the mean are provided. Neuroreceptor drive network consistency was estimated by correlating the estimated drive networks from the HCP, LSD, Modafinil, and UCLA5c datasets.

## Data Availability

Minimally pre-processed Human Connectome Project data can be downloaded from https://www.humanconnectome.org/. Neuroreceptor composition data can be obtained at https://github.com/netneurolab/hansen_receptors. LSD dataset can be obtained at https://openneuro.org/datasets/ds003059/versions/1.0.0. Modafinil dataset can be obtained at https://openneuro.org/datasets/ds000133/versions/00001. UCLA Consortium for Neuropsychiatric Phenomics Dataset can be downloaded at https://openfmri.org/dataset/ds000030/. The brain connectivity toolbox can be downloaded at https://sites.google.com/site/bctnet/. Cortical geometric eigenmode data can be obtained at https://osf.io/xczmp/.

## Acknowledgements

We would like to thank Javier Garcia, Jessie J Polanco and Italo’Ivo Lima Dias Pinto for their thoughtful comments. This research was supported by the U.S. Army DEVCOM Army Research Laboratory through army educational outreach program (W911SR-15-2-0001). The views and conclusions contained in this document are those of the authors and should not be interpreted as representing the official policies, either expressed or implied, of the U.S. Army DEVCOM Army Research Laboratory or the U.S. Government. The U.S. Government is authorized to reproduce and distribute reprints for Government purposes notwithstanding any copyright notation herein.

## Author Contributions

Conceptualization: JN

Methodology: JN

Data Curation: JN

Visualization: JN

Funding acquisition: JN, KB

Supervision: KB

Writing – original draft: JN, KB

Writing – review & editing: JN, KB

