## Supplemental Information for "The neuroreceptors and transporters underlying spontaneous brain activity"

Neuroreceptors associated with altered BOLD signal in Schizophrenia

In schizophrenia the current dominant hypothesis suggests a dysregulation in the dopamine system – Dopamine Hypothesis^1^. As a result currently, most licensed treatments focus on dopamine by blocking the D2/D3 neuroreceptors^2^. However, treatments focusing on dopamine show limited efficacy particularly for negative symptoms^3^. The limitations of the dopamine hypothesis and of medication targeting dopamine suggests that other neurotransmitter systems might play a role.

Other work suggests that schizophrenia may be linked to NMDA, NET, 5-HT1a, 5-HT4, GABAa and α4β2 neuroreceptors. One alternative hypothesis suggests that hypofunction of the NMDA glutamate neuroreceptor may be the cause of the illness^4^. The role of NMDA is supported by observations that NMDA antagonist drugs such as ketamine and phencyclidine produce features that mimic those observed in schizophrenia^5^. Moreover, copy number variant and GWAS studies have implicated genes that encode NMDA^6^. Additionally, cognitive symptoms may be explained by abnormal functioning of the locus coeruleus which produces norepinephrine^7^. This is supported by both pharmacological^8–11^ and animal models^12^ which suggest that norepinephrine plays a critical role in schizophrenia. Moreover, serotonin is also thought to be associated with schizophrenia and is supported by NMR spectroscopy and PET imaging^13^. Note that the analysis identified NMDA, norepinephrine transporter (NET), serotonin 5-HT1a and 5-HT4 to be associated with altered brain activity in the schizophrenia aligning with this prior work. Studies have also implicated GABA to play a role in schizophrenia^14–16^. Additionally, the analysis identified an association between nicotinic acetylcholine receptor (α4β2) and schizophrenia aligning with prior work^17^. In fact, the newest medication approved for treating schizophrenia in over 30 years, KarXT, targets the cholinergic neurotransmitter system^18^. Overall, the results from the analysis in Figure 5A align with the alternative hypotheses to dopamine in schizophrenia.

Along these lines, prior work on psychosis-spectrum disorders that leveraged AHBA gene expression data and fMRI identified associations between functional connectivity activity and serotonin and GABA^19^. This prior work aligns in part with our findings reflecting a relationship between the serotonin 5-HT1a and 5-HT4 neuroreceptors, and a trending relationship with GABAa. Moreover, we identified associations between NET, NMDA, and α4β2 neuroreceptors and brain activity. This divergence between our findings are those of Ji et. al. 2021 could reflect the dissociation between mRNA levels and neuroreceptor density^20^. Additionally, prior work by Lawn et. al., 2024 used neuroreceptor maps to link them to altered brain activity in the UCLA5 dataset^21^. Using a dual regression analysis – REACT – the findings in Lawn et. al., 2024 implicate VAChT (cholinergic) neuroreceptor in schizophrenia, the mGLuR5 (glutamate) and GABAa neuroreceptors in both schizophrenia and bipolar disorder. On the one hand, the specific neuroreceptors identified by Lawn et. al., 2024 do not align with what we have identified, on the other hand, the general class of neurotransmitter systems do align – i.e., cholinergic, glutamate, and GABA. It is important to note that, Lawn et. al., 2024 limited their investigation to NET, DAT, 5-HTT, VAChT, mGLuR5, and GABAa. However, the one difference is that our analysis identified NET unlike in Lawn et. al., 2024 this could reflect processing difference. We were more aggressive in removing high motion volumes. Additionally, REACT is a dual regression model in which explainable variance might be diminished with each regression step.

It is important to note that the individuals in the schizophrenia group are medicated with dopamine modulating drugs. As a result, the medication could be moderating the effects of dopamine and therefore dampening the association with schizophrenia. Therefore, caution is warranted when interpreting these results given the small sample size and since there is no ground truth in the molecular factors underlying these disorders.

Neuroreceptors associated with altered BOLD signal in ADHD

Attention has been shown to be associated with acetylcholine, dopamine, serotonin and norepinephrine^22^. Clinical studies suggest that stimulating nicotinic cholinergic receptors can alleviate symptoms and improve executive function in individuals with ADHD^23–25^. Moreover, adults with ADHD exhibit a reduction in symptoms when administrated an α4β2 nicotinic agonists^26^. In line with this prior work our analysis identified an association between α4β2 and altered brain activity in individuals with ADHD. Additionally, hypoactivity in the dopaminergic pathway is considered to play a role in the deficits seen in ADHD patients^27–29^. In line with this prior work, we observed a significant association with the dopamine D1 neuroreceptor. Besides the association between the D1 and α4β2, our results indicated at trending association between the serotonin 5-HT4 neuroreceptor and brain activity in ADHD. The evidence for a relationship with between serotonin and ADHD is mixed, but medication that act on serotonin system are second-line drug of choice for treating ADHD^30^. Additionally, we observed an association between alterations in brain activity in the ADHD group with GABAa neuroreceptor. This observation is in line with previous work showing alterations in GABA levels in children with ADHD^31,32^. Interestingly, we did not observe a relationship between norepinephrine transporter (NET) and ADHD. In particular, ADHD has been associated with decreased NET density^33^. This dissociation with prior work could reflect effects of the medication the individuals are administered. Additionally, the observed differences, for instance in functional connectivity, between healthy individuals and those with ADHD were subtle suggesting that the analysis may not as be sensitive.

Comparing Models: neuroreceptors vs. structural and geometric modes

The main focus of our analysis was on the relationship between neuroreceptors, transporters and the BOLD activity, but the BOLD signal can be reconstructed from anatomical factors – structural connectivity^34,35^ and cortical folding geometry^36^. It is an open question as to which model – neuroreceptor, structural or geometric – accurately and parsimoniously reconstructs spontaneous brain activity.

Toward this end, we compared the accuracy of the neuroreceptor-based reconstructed maps of the dominant spatial patterns of the BOLD signal with reconstructed maps obtained using structural and geometric modes. For this analysis, we utilized the BOLD signal from the HCP dataset due to its extensive scanning duration. Specifically, the BOLD signal was concatenated across participants resulting in 200 x 57,600 data points. The neuroreceptor-based framework reconstructed BOLD components with an accuracy of R > 0.64, significantly exceeding both null models (P_spatial_ < 0.001; P_spin_ < 0.001; **Fig. S12**). Comparing models, the neuroreceptor-based reconstruction was more parsimonious for 8 out of the 10 BOLD components (**Fig. S13**). Only components C4 and C6 required fewer GC modes than the neuroreceptor analysis. However, these components accounted for only 4.01% and 2.57% of the spatiotemporal variance in the BOLD signal, respectively.

Communication between brain regions and maintaining healthy brain states is reliant on anatomical factors and anatomical changes are associated with neuropathology, highlighting the importance of anatomical factors^37,38^. These results suggest that a neuroreceptor-based framework may be better at estimating brain activity since the concentration of neuroreceptors on dendritic spines can amplify signals from weak structural connections or diminish signals from strong physical connections estimated from diffusion MRI^39^. Additionally, brain regions that share similar neuroreceptor and transporter profiles are more likely to exhibit similar patterns in brain activity since they would presumably respond to the same neuromodulators.

It is important to note that the different spatial resolutions, smoothing levels, levels of autocorrelation, and scanner of origin associated with the neuroreceptor maps may account for why the neuroreceptor-based framework is more parsimonious than models based on structural connectivity and cortical folding geometry. Future studies could minimize these concerns by harmonizing the scanning parameters and/or pre-processing of PET imaging.

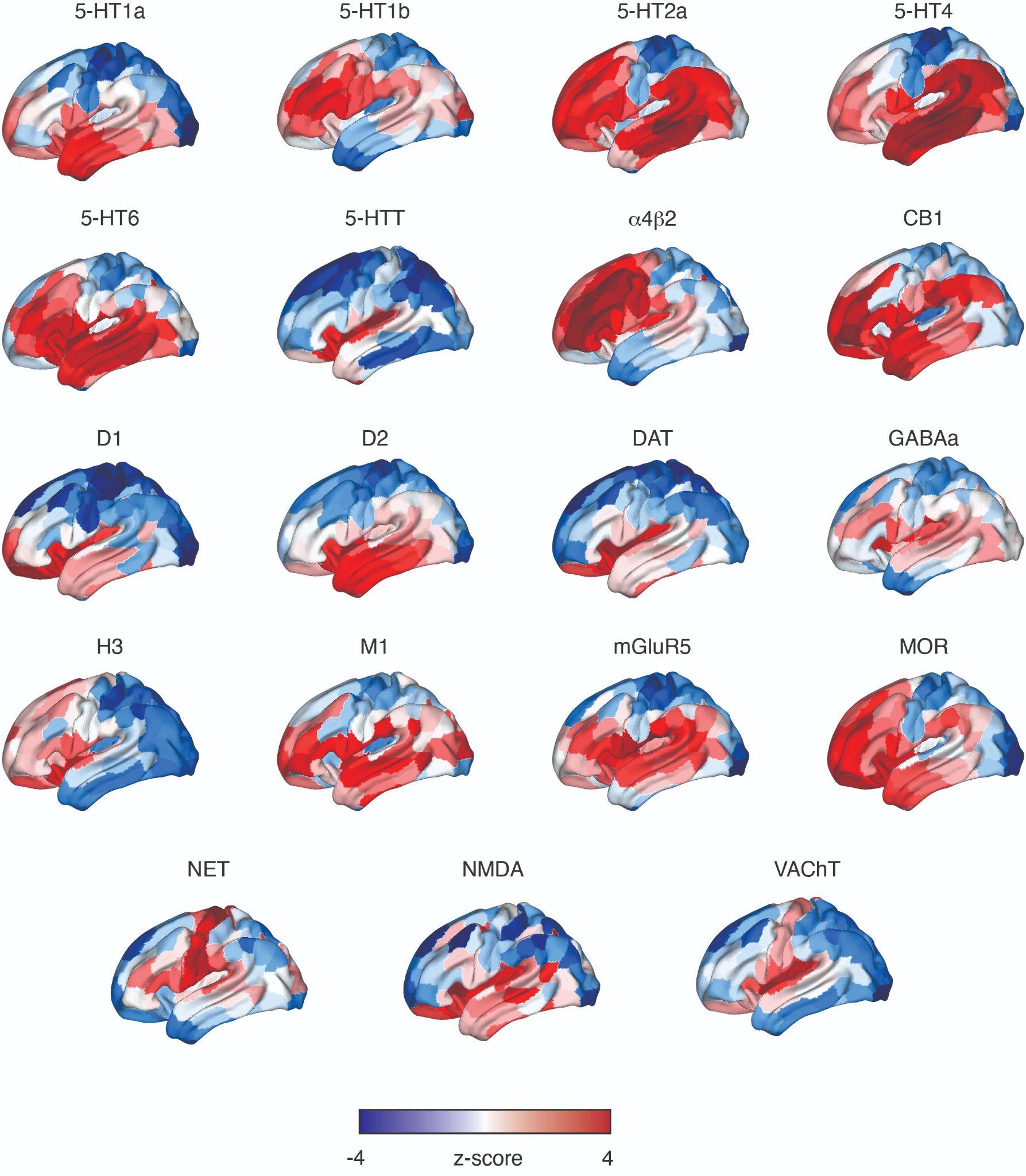

**Figure S1**. **Neuroreceptor and transporter density spatial maps.** Density maps of 19 neuroreceptors and transporters across the cortex from Hansen *et. al.*, 2022^40^. The density for each neuroreceptor and transporter was estimated in 200 brain regions part of the Schaefer atlas by averaging all voxels within an ROI.

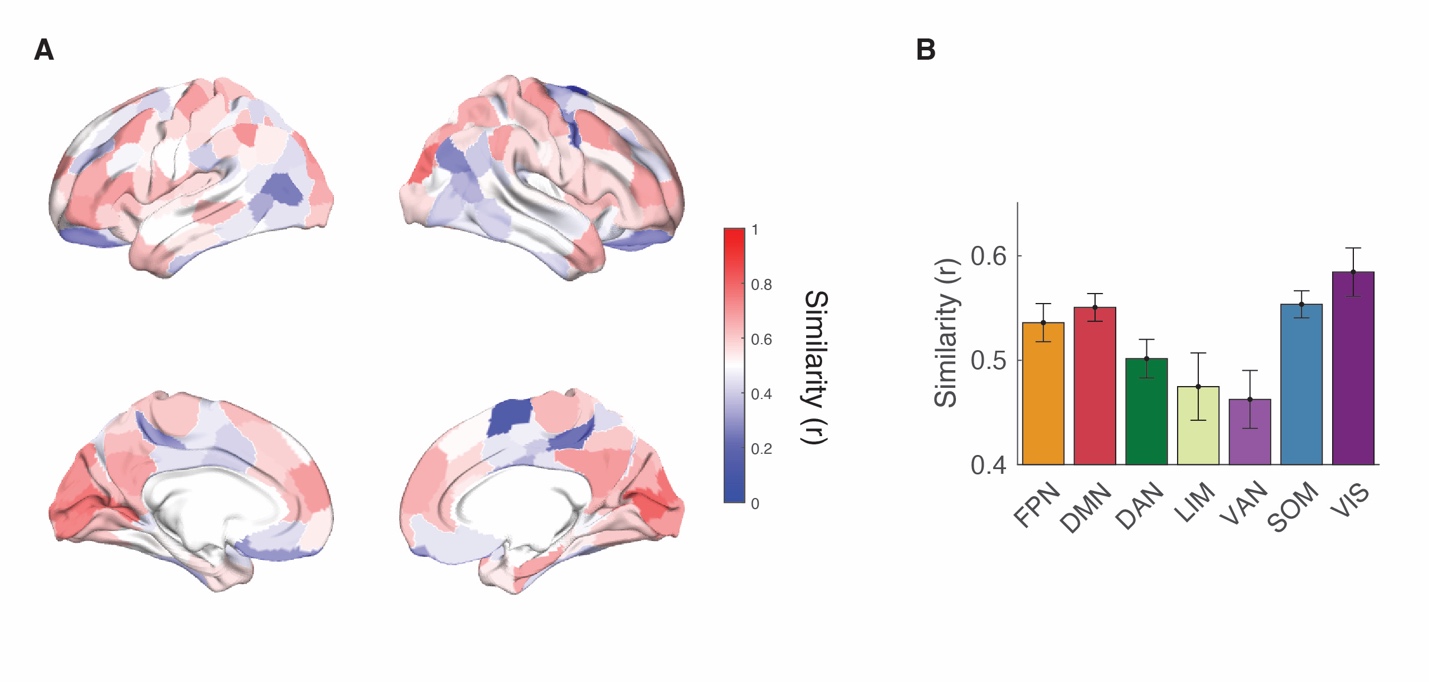

**Figure S2**. **Brain region and network level reconstruction accuracy.** (A) Reconstruction accuracy for each of the 200 brain regions part of the Schaefer atlas. Accuracy was estimated at the correlation between empirical and reconstructed BOLD signal concatenated across subjects. B) Average reconstruction accuracy across brain network part of the Schaefer atlas. Error bars depict Mean ± SEM. FPN, Frontal Parietal Network; DMN, Default Mode Network; DAN, Dorsal Attention Network; LIM, Limbic Network; VAN, Ventral Attention Network; SOM, Somatomotor Network; VIS, Visual Network.

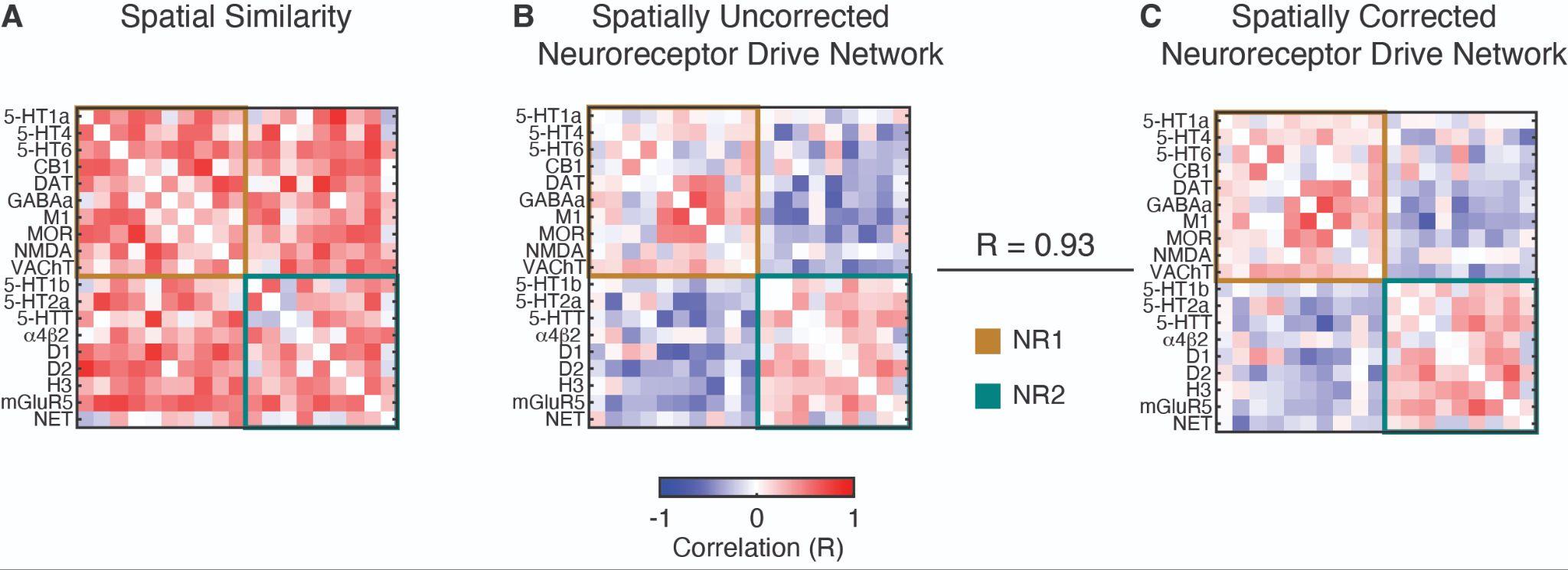

**Figure S3. Relationship between neuroreceptor drive network and spatial similarity.** (A) The similarity in density maps among the 19 neuroreceptors and transporters. Spatial similarity was estimated using the Pearson correlation between neuroreceptor maps. The spatial similarity network has been organized to reflect the two modules – NR1 and NR2 – identified in the main analysis. (B) Neuroreceptor drive network, same as Fig. 2 panel B. (C) Spatially corrected neuroreceptor drive network. The spatial similarity was regressed out from the drive network in plane B.

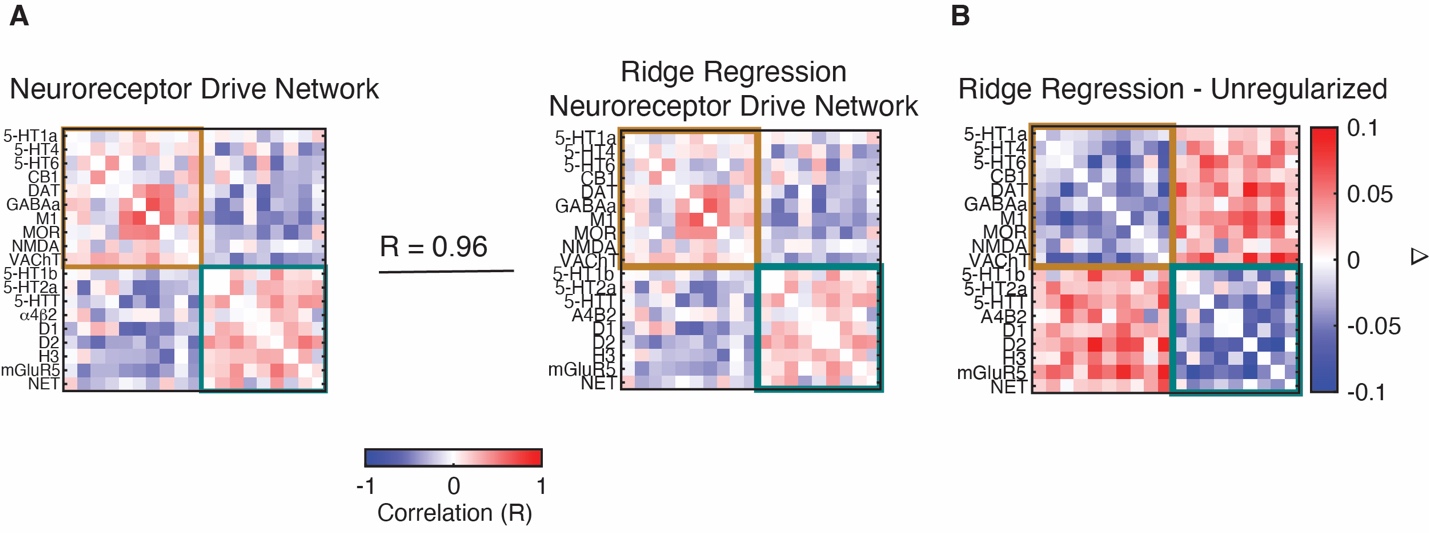

**Figure S4**. **Relationship between unregularized and ridge regression regularized β-values.** (A) Comparison between the unregularized neuroreceptor drive network (*left*), same as Fig. 2 panel B, with beta-values estimated using ridge regression (*right*). (B) Differences in neuroreceptor drive network values between unregularized and ridge regression.

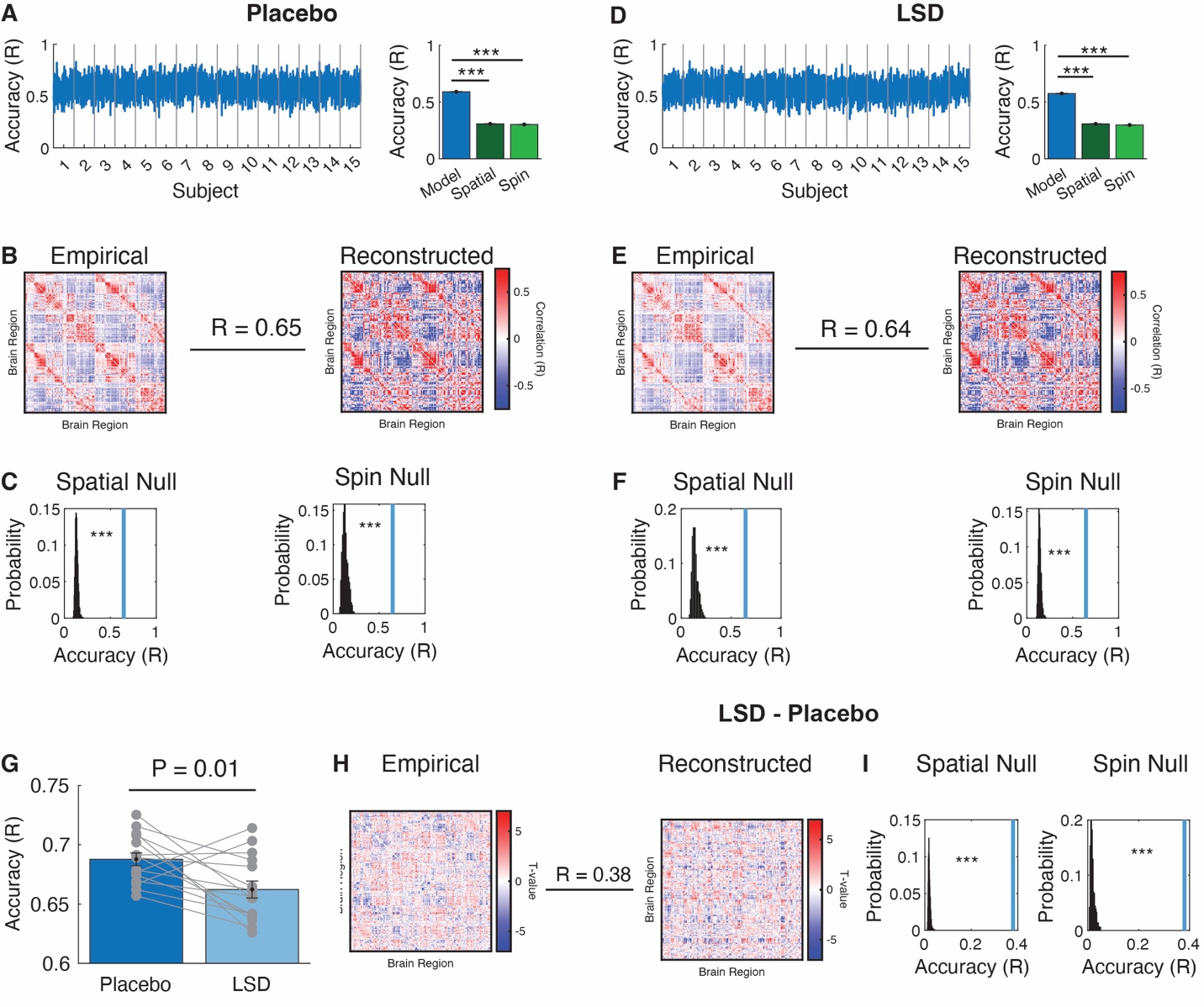

**Figure S5. Reconstruction of spontaneous brain activity before administration of LSD.** (A) Model accuracy of the BOLD signal reconstructed for each period of time after the administration of the placebo (*left*). Note that temporal data reflects all timepoints pooled across the 15 participants (434 data points per participant). Neuroreceptor-based model performance compared to spatial and spin null models (*right*). Bar plots show the average accuracy at the group-level and for the spatial and spin null models across 1000 iterations. Error bars show mean ± sem. Statistical significance was estimated using an independent sample t-test. (B) Accuracy between the empirical and reconstructed functional connectivity after the administration of the placebo. (C) Similarity in functional connectivity is significantly greater than both the permutation null model and spin null model. Blue line depicts similarity value from panel B. (D-F) Sam as plane A, B and C, but for LSD. (G) Difference in reconstruction accuracy of the BOLD signal between the placebo and LSD conditions. Error bars depict mean ± sem. Accuracy differences were estimated using paired-samples t-test. Accuracy values from panels A and D were first Fisher r-to-z transformed and averaged within each participant, respectively. (H) Similarity in estimated differences in functional connectivity between empirical and reconstructed. (I) Observed similarity from panel H is significantly greater than permuted and spin null models. Blue line depicts similarity value from panel H. *** P < 0.001.

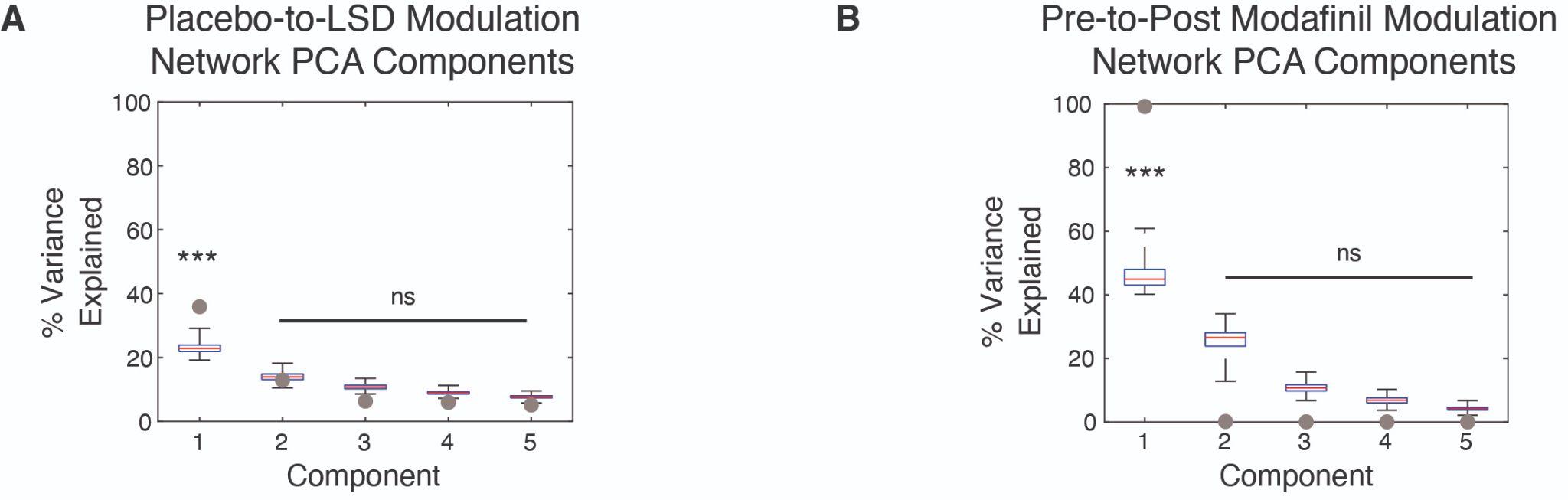

**Figure S6. Variance explained by each component of the modulation network for the LSD and Modafinil.** (A) The variance explained by the top 5 PCA components of the modulatory network to transition the drive network from the placebo to the LSD state. (B) Same as panel A, but for Modafinil. Gray dots depict the observed amount of variance in the modulation network explained by each principal component. Box plots represent the variance explained across 1000 permuted modulation networks. ***P_perm_ < 0.001; ns, not significant.

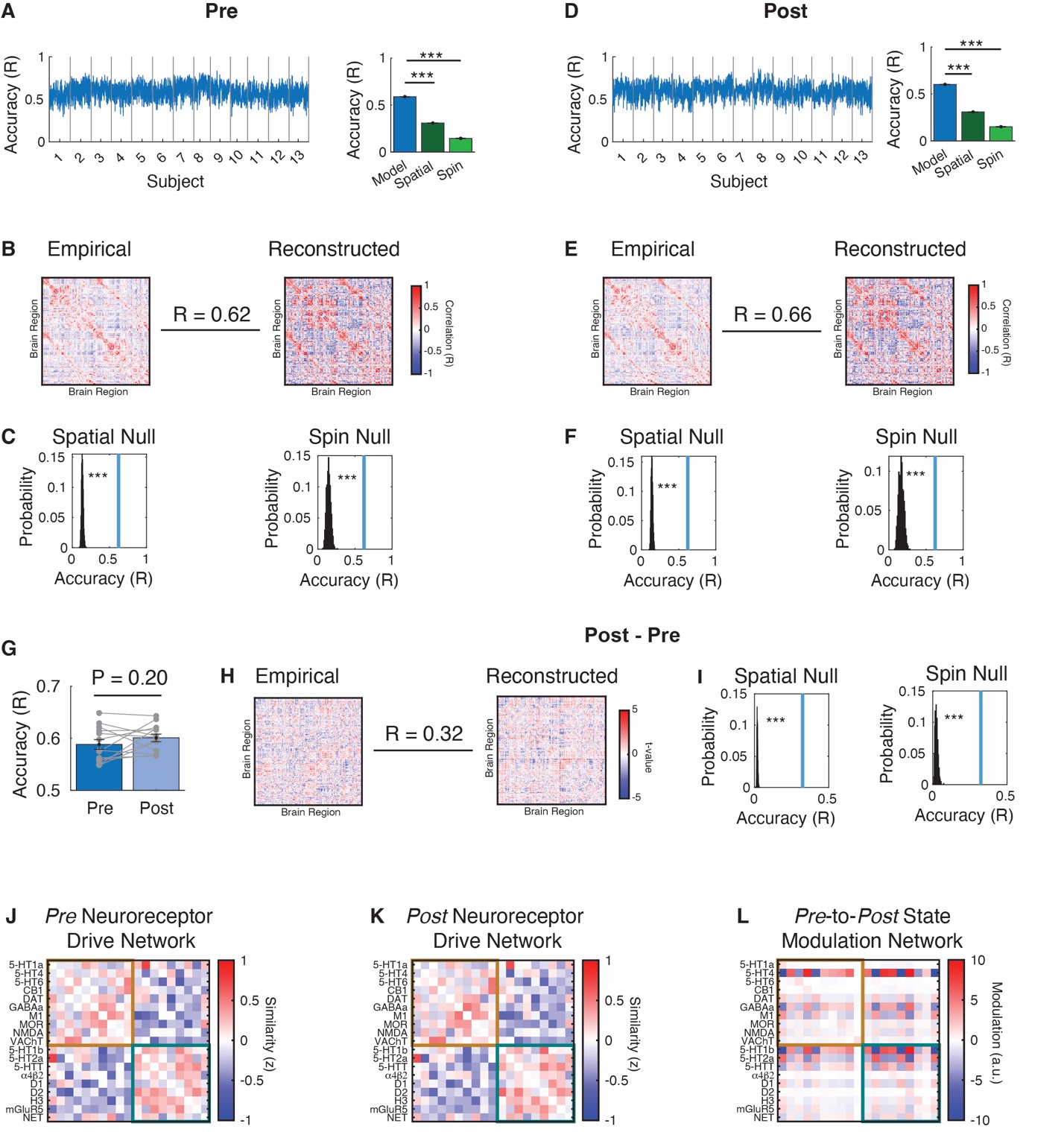

**Figure S7. Reconstruction of spontaneous brain activity *pre* and *post* administration of Modafinil.** (A) Model accuracy of the BOLD signal reconstructed for each period of time *pre* administration of Modafinil (*left*). Neuroreceptor-based model performance compared to spatial and spin null models (*right*). Bar plots show the average accuracy at the group-level and for the spatial and spin null models across 1000 iterations. Error bars show mean ± sem. Statistical significance was estimated using an independent sample t-test. (B) Accuracy between the empirical and reconstructed functional connectivity *pre* administration of Modafinil (*left*). Neuroreceptor-based model performance compared to spatial and spin null models (*right*). Bar plots show the average accuracy at the group-level and for the spatial and spin null models across 1000 iterations (*right*). (C) Similarity in functional connectivity is significantly greater than both the permutation null model and spin null model. Blue line depicts similarity value from panel B. (D-F) Sam as plane A, B and C, but for *post* administration of Modafinil. (G) Difference in reconstruction accuracy of the BOLD signal *pre* and *post* administration of Modafinil. Error bars depict mean ± sem. Accuracy differences were estimated using paired-samples t-test. Accuracy values from panels A and D were first Fisher r-to-z transformed and averaged within each participant, respectively. (H) Similarity in estimated differences in functional connectivity between empirical and reconstructed. (I) Observed similarity from panel H is significantly greater than permuted and spin null models. Blue line depicts similarity value from panel H. (J) Group-level neuroreceptor drive network *pre* administration of Modafinil estimated from the drive values from each of the 19 neuroreceptor and transporter using Pearson correlation. The network is organized according to the module structure identified in the HCP dataset. (K) Group-level neuroreceptor drive network *post* administration of Modafinil, structured similarly to panel A. (L) Modulation network illustrating shifts in brain activity from *pre* to *post* Modafinil induced state. The modulation network is derived by multiplying the inverse of the *pre* neuroreceptor drive network (Panel J) with the *post* drive network (Panel K). ***P < 0.001

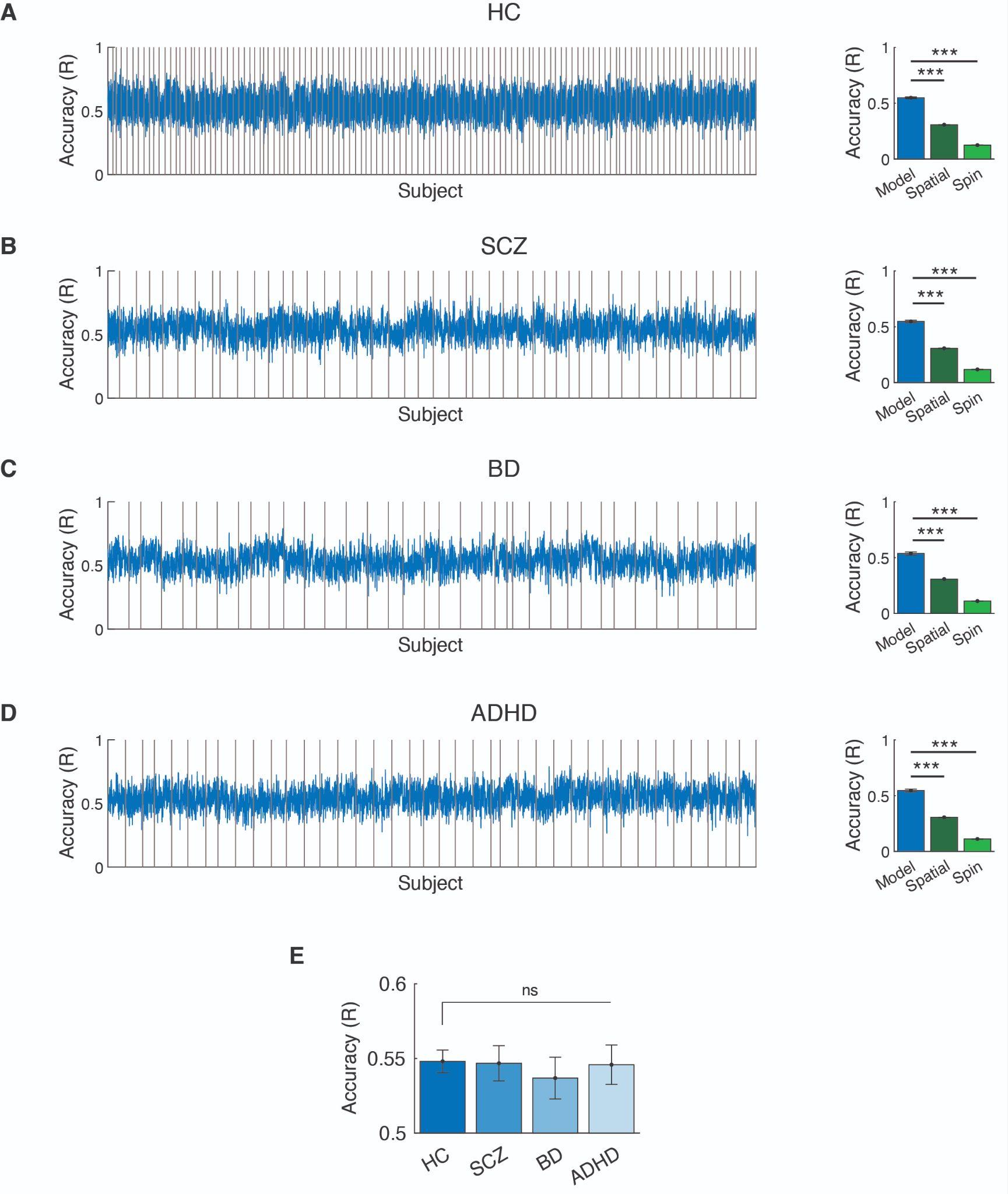

**Figure S8. BOLD signal reconstruction accuracy for the LA5c dataset.** (A) BOLD signal reconstruction accuracy for healthy controls (HC; *left*). Vertical lines separate participants. Neuroreceptor-based model performance compared to spatial and spin null models (*right*). Bar plots show the average accuracy at the group-level and for the spatial and spin null models across 1000 iterations. Error bars show mean ± sem. Statistical significance was estimated using an independent sample t-test. (B-D) same as panel A, but for schizophrenia (SCZ), bipolar disorder (BD), and ADHD, respectively. (E) Difference in reconstruction accuracy of the BOLD signal between the healthy control group and schizophrenia, bipolar disorder and ADHD group. Differences in reconstruction accuracy were evaluated using independent samples t-test between the groups. Error bars depict mean ± sem. ns, not significant.

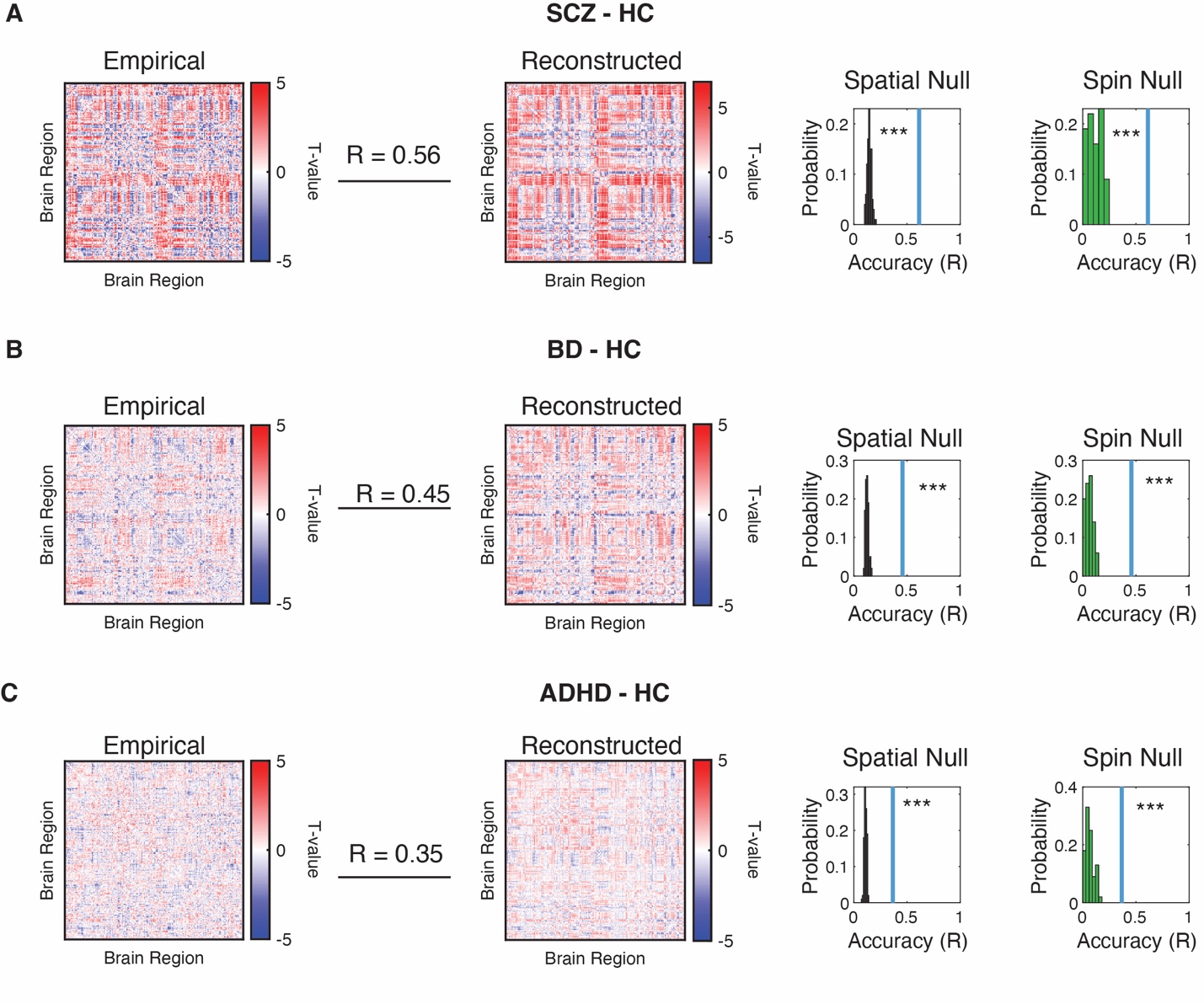

**Figure S9. Similarity in estimated differences in functional connectivity between neuropsychiatric and healthy controls.** (A) Similarity between empirical differences in functional connectivity between schizophrenia (SCZ) and healthy controls and reconstructed FC maps*.* Observed similarity was significantly greater than both the permutation and spin null models. (B,C) Same as panel A, but for bipolar disorder (BD) and ADHD. ***P < 0.001.

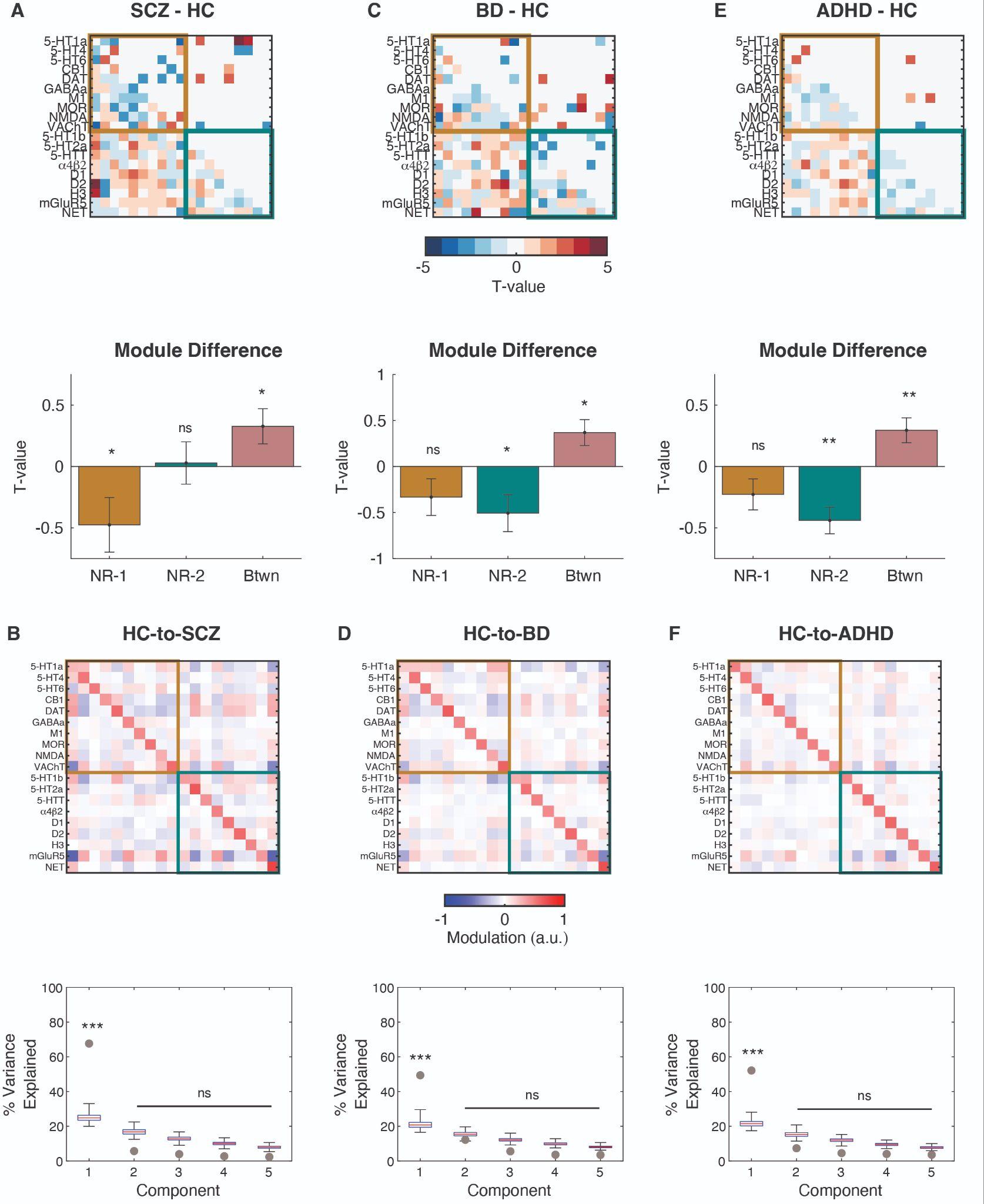

**Figure S10. Schizophrenia, bipolar disorder and ADHD associated neuroreceptor drive networks and modulation networks.** (A) Schizophrenia (SCZ) induces changes in the neuroreceptor drive network. The top panel shows differences in the drive network between schizophrenia patients and healthy controls (HC), assessed using paired samples t-tests. R-values were Fisher z-transformed before statistical testing. The top half of the network represents statistically significant changes (independent samples t-test; P < 0.05, uncorrected), while the bottom half shows the magnitude of all pairwise changes. The bottom panel displays changes in connectivity within and between modules, with drive networks organized according to the module structure identified in the HCP dataset. A one-sample t-test was used to evaluate deviations from zero. (B) Modulation network that shifts the brain activity from brain state associated with healthy controls to schizophrenia (top). Bottom panel shows the variance explained by the top 5 PCA components of the modulatory network to transition the drive network from the brain state associated with healthy controls to schizophrenia. Circle depicts the observed amount variance in the modulation network explained by a principal component. Box plots represent the variance explained across 1000 permuted modulation networks. (C,D) Same as panel A and B, but for bipolar disorder, respectively. (E,F) Same as panel A and B, but for ADHD, respectively. ***P_perm_ < 0.001; ns, not significant.

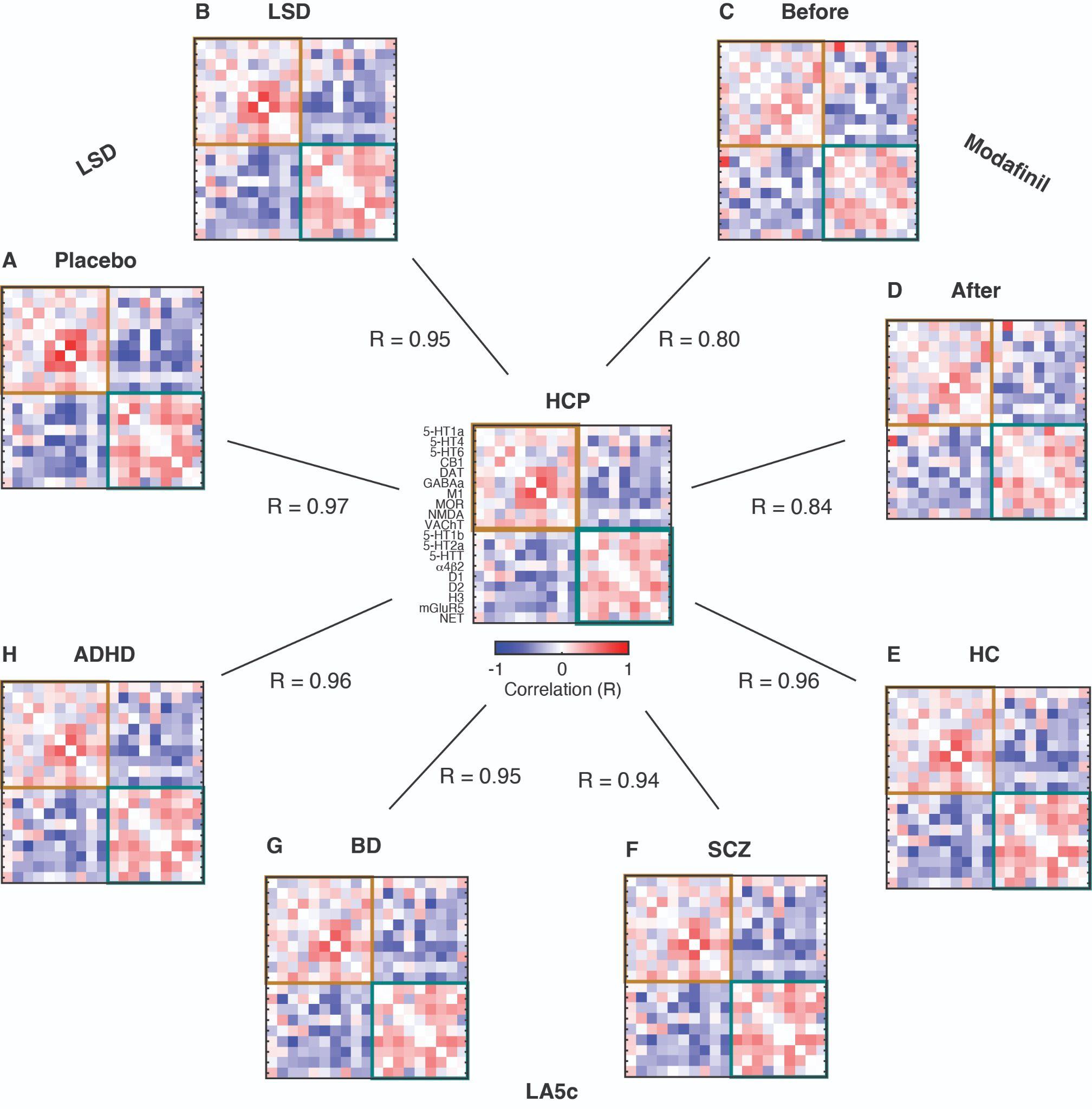

**Figure S11. Similarity of neuroreceptor drive networks across datasets.** Group-level similarity of neuroreceptor drive networks estimated from the HCP dataset compared with: LSD dataset (A) Placebo condition, (B) LSD condition; Modafinil dataset (C) Before and (D) After; and LA5c dataset (E) Healthy Controls (HC), (F) Schizophrenia (SCZ), (G) Bipolar Disorder (BD), and (H) ADHD. Drive networks are oriented according to the modular organization identified in the HCP dataset.

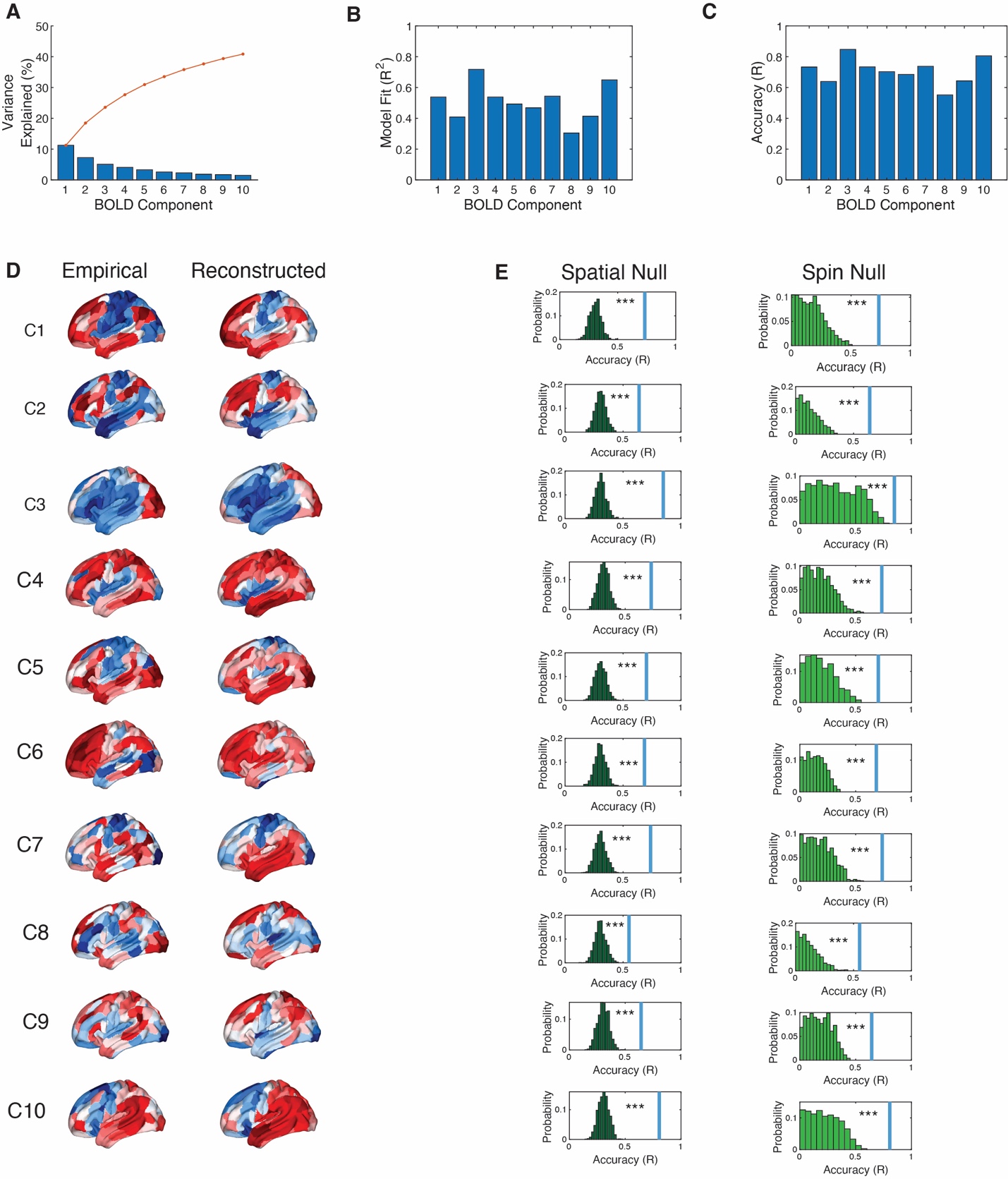

**Figure S12**. **Modeling the BOLD signal components.** (A) The first 10 principal components of the fMRI BOLD signal account for 40.86% of the signal. (B) The variance explained by the neuroreceptor model (R^2^) for each BOLD signal component. (C) The accuracy of the neuroreceptor model when reconstructing BOLD components. Accuracy is estimated using the Pearson correlation between the empirical and reconstructed components. (D) Brain maps of the empirical (*left*) and reconstructed (*right*) BOLD components C1 to C10. (E) The accuracy from the neuroreceptor model is significantly greater than the accuracy obtained using the permutation and spin null models. Blue line represents the accuracy from the neuroreceptor model. Histograms depict the distribution in accuracy obtained from 1000 randomization in the permutation and spin null models.

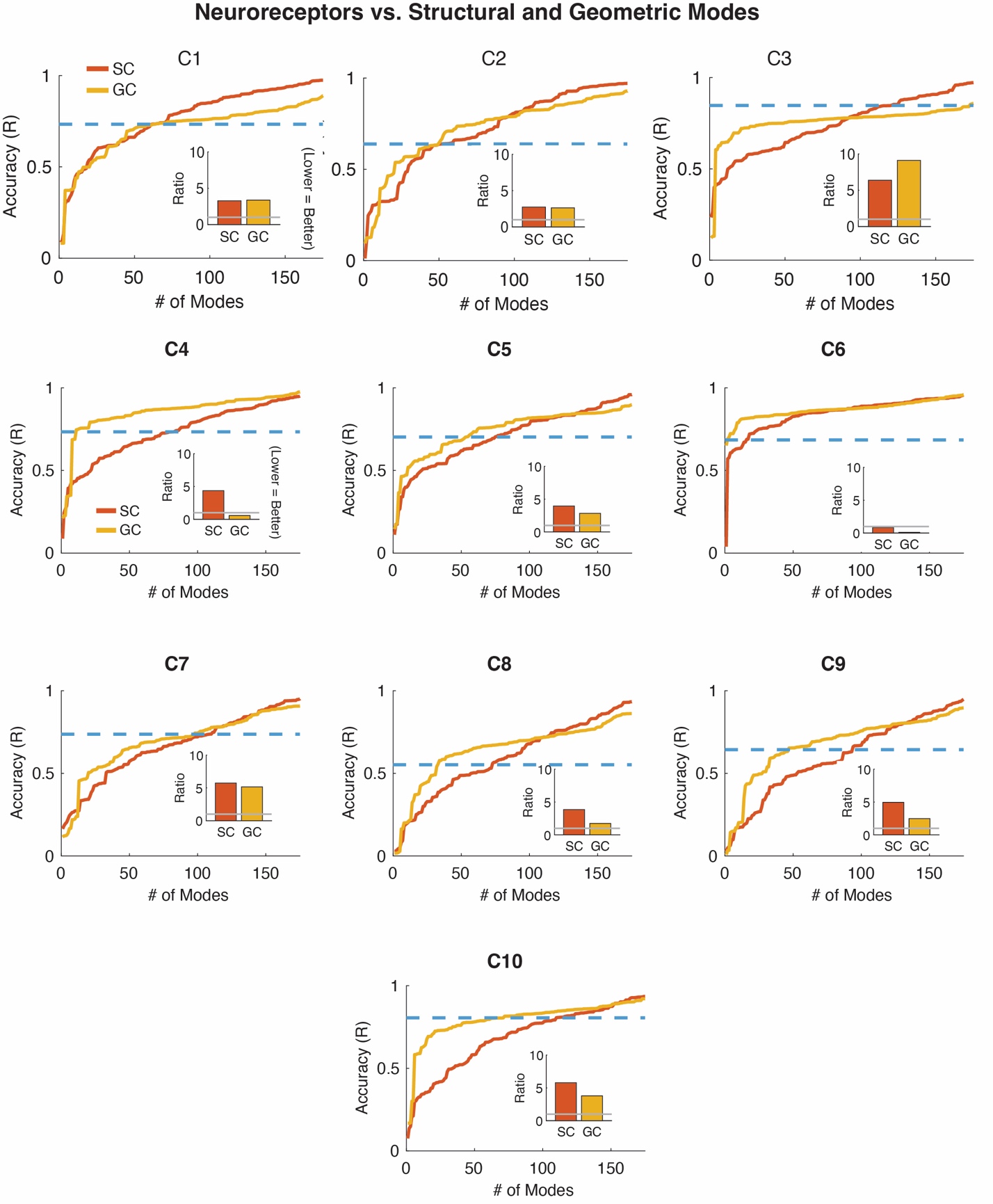

**Figure S13. The number of geometric and structural modes required to reconstruct BOLD components C1-C10 with the same accuracy as the neuroreceptor model.** The number geometric or structural modes required to achieve the same level of accuracy in the PCA component C1-C10 of the BOLD signal as the neuroreceptor model. The geometric or structural modes require at least twice as many modes as the neuroreceptor model. The exceptions are C4 and C6 in which the geometric model required 11 and 2 modes for the model to achieve the same accuracy compared to the 19 in the neuroreceptor model, respectively. However, BOLD components C4 and C6 account for only 4.01% and 2.57% of the variance in the BOLD signal.
